# Symmetry and simplicity spontaneously emerge from the algorithmic nature of evolution

**DOI:** 10.1101/2021.07.28.454038

**Authors:** Iain G Johnston, Kamaludin Dingle, Sam F. Greenbury, Chico Q. Camargo, Jonathan P. K. Doye, Sebastian E. Ahnert, Ard A. Louis

## Abstract

Engineers routinely design systems to be modular and symmetric in order to increase robustness to perturbations and to facilitate alterations at a later date. Biological structures also frequently exhibit modularity and symmetry, but the origin of such trends is much less well understood. It can be tempting to assume – by analogy to engineering design – that symmetry and modularity arise from natural selection. But evolution, unlike engineers, cannot plan ahead, and so these traits must also afford some immediate selective advantage which is hard to reconcile with the breadth of systems where symmetry is observed. Here we introduce an alternative non-adaptive hypothesis based on an algorithmic picture of evolution. It suggests that symmetric structures preferentially arise not just due to natural selection, but also because they require less specific information to encode, and are therefore much more likely to appear as phenotypic variation through random mutations. Arguments from algorithmic information theory can formalise this intuition, leading to the prediction that many genotype-phenotype maps are exponentially biased towards phenotypes with low descriptional complexity. A preference for symmetry is a special case of this bias towards compressible descriptions. We test these predictions with extensive biological data, showing that that protein complexes, RNA secondary structures, and a model gene-regulatory network all exhibit the expected exponential bias towards simpler (and more symmetric) phenotypes. Lower descriptional complexity also correlates with higher mutational robustness, which may aid the evolution of complex modular assemblies of multiple components.

---

Evolution proceeds through genetic mutations which generate the novel phenotypic variation upon which natural selection can act. The relationship between the space of genotypes and the space of phenotypes can be encapsulated as a genotype-phenotype (GP) map [1–3]. These can be viewed algorithmically, where random genetic mutations search in the space of (developmental) algorithms encoded by the GP map, a relationship that has been highlighted, for example, in plants [4], in Dawkins’ ‘biomorphs’ [5] and in molecules [6].

Genetic mutations are random in the sense that they occur independently of the phenotypic variation they produce. This does not, however, mean that the probability *P*(*p*) that a GP map produces a phenotype *p* upon random sampling of genotypes will be anything like a uniformly random distribution. Instead, highly general (but rather abstract) arguments based on the coding theorem of algorithmic information theory (AIT) [7], predict that the *P*(*p*) of many GP maps should be highly biased towards phenotypes with low Kolmogorov complexity *K*(*p*) [8]. High symmetry can, in turn, be linked to low *K*(*p*) [6, 9–11]. An intuitive explanation for this algoritmic bias towards symmetry proceeds in two steps: 1.) Symmetric phenotypes typically need less information to encode algorithmically, due to repetition of subunits. This higher compressibility reduces constraints on genotypes, implying that more genotypes will map to simpler, more symmetric phenotypes than to more complex asymmetric ones [2, 3]. 2.) Upon random mutations these symmetric phenotypes are much more likely to arise as potential variation [12, 13], so that a strong bias towards symmetry may emerge even without natural selection for symmetry.

## Symmetry in protein quaternary structure and polyominoes

We first explore evidence for this algorithmic hypothesis by studying protein quaternary structure, which describes the multimeric complexes into which many proteins self-assemble in order to perform key cellular functions (Fig. 1A and Supporting Information (SI) Fig S1 and section S1). These complexes can form in the cell if proteins evolve attractive interfaces allowing them to bind to each other [14–16]. We analysed a curated set of 34,287 protein complexes extracted from the Protein Data Bank (PDB) that were categorised into 120 different bonding topologies [16]. In Fig. 1B, we plot, for all complexes involving 6 subunits (6-mers), the frequency with which a protein complex of topology *p* appears against the descriptional complexity 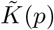, an approximate measure of its true Kolmogorov assembly complexity *K*(*p*), defined here as the minimal number of distinct interfaces required to assemble the given structure under general self-assembly rules (Methods). Here 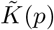 can also be thought of as a measure of the minimal number of evolutionary innovations needed to make a self-assembling complex. The highest probability structures all have relatively low 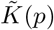. Since structures with higher symmetry need less information to describe [6, 9–11], the most frequently observed complexes are also highly symmetric. Figs. 1C and Figs S2A & S3A further demonstrate that structures found in the PDB are significantly more symmetric than the set of all possible 6-mers (Methods). Similar biases towards high symmetry structures obtain for other sizes (Fig S2B).

**FIG. 1.**
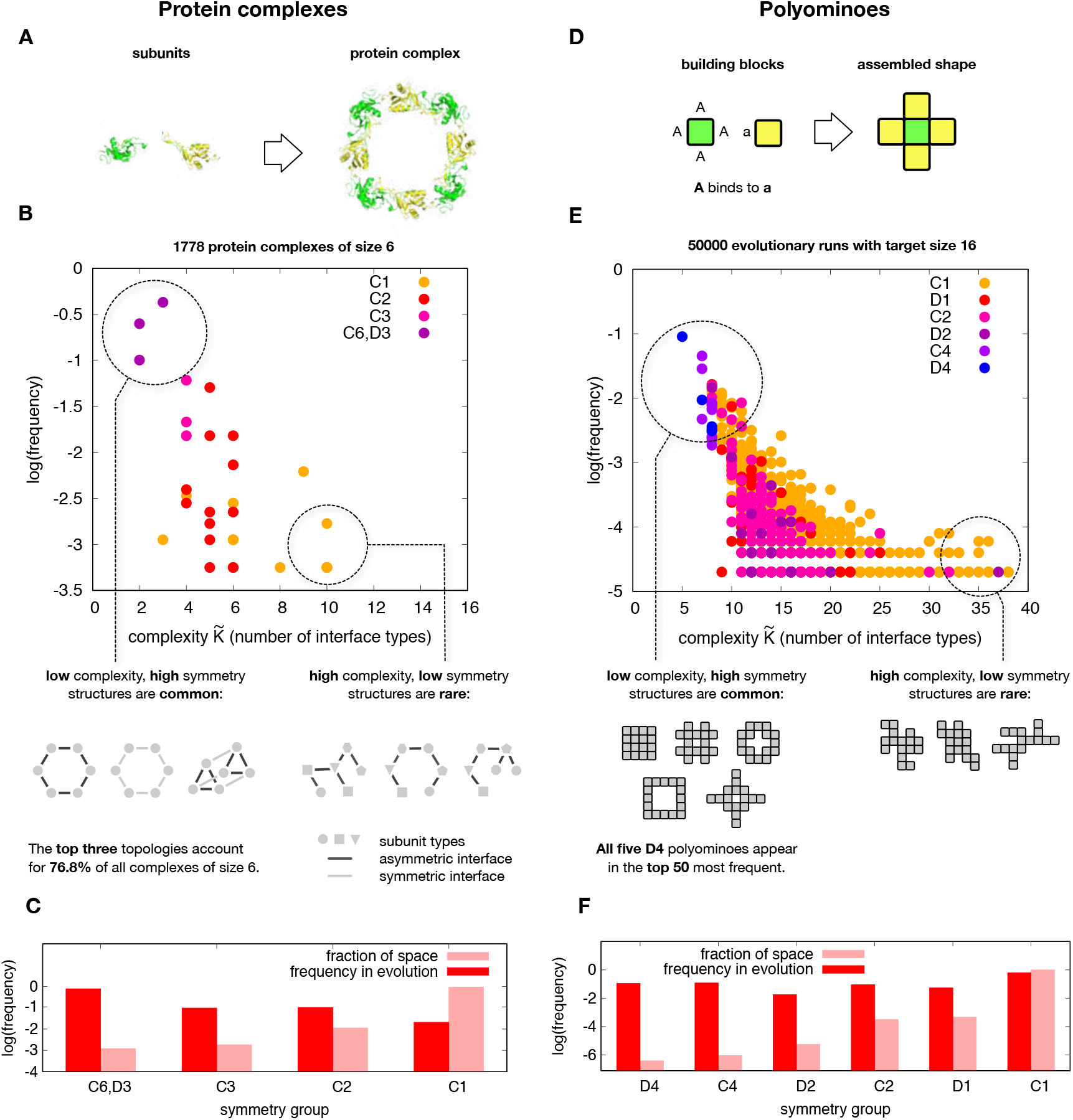
(A) Protein complexes self-assemble from individual units. (B) Frequency of 6-mer protein-complex topologies found in the PDB versus the number of interface types, a measure of complexity 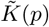. Symmetry groups are in standard Schoenflies notation: *C*_6_, *D*_3_, *C*_3_, *C*_2_, *C*_1_. There is a strong preference for low complexity/high symmetry structures. (C) Histograms of the frequencies of symmetries for 6-mer topologies found in the PDB (dark red) versus the frequencies by symmetry of the morphospace of all possible 6-mers illustrate that symmetric structures are hugely over-represented in the PDB database. (D) Polyomino complexes (here *a* binds to *A*) self-assemble from individual units just as the proteins do. (E) The frequency of polyominoes that fix in evolutionary simulations with a fitness maximum at 16-mers versus the number of interface types (a measure of complexity 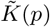) exhibits a strong bias towards high symmetry structures, similar to protein complexes. (F) Histograms of the frequency of symmetry groups for all 16-ominoes (light) and for 16-ominoes appearing in the evolutionary runs (dark), quantify how strongly biased variation drives a pronounced preference for high symmetry structures.

In order to understand the evolutionary origins of this bias towards symmetry we turn to a tractable GP map for protein quaternary structure. In the Polyomino GP map, two-dimensional tiles self-assemble into polyomino structures [17] that model protein-complex topologies [18] (Fig. 1D). The sides represent the interfaces that bind proteins together. Within the Polyomino GP map, the genomes are bit strings used to describe a set of the tiles and their interactions. The phenotypes are polyomino shapes *p* that emerge from the self-assembly process. Although this model is highly simplified, it has successfully explained evolutionary trends in protein quaternary structure such as the preference of dihedral over cyclic symmetry in homomeric tetramers [15, 17], or the propensity of proteins to form larger aggregates such as haemoglobin aggregation in sickle-cell anaemia [18].

To explore the strong preference for simple structures, we performed evolutionary simulations where fitness is maximised for polyominoes made of 16 blocks (Methods). With 16 tile types and 64 interface types, the GP map denoted as 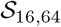 allows all 13,079,255 possible 16-mer polyomino topologies (SI Table I) to be made. Fig 1E demonstrates that evolutionary outcomes are exponentially biased towards 16-mer structures with low 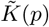 (using the same complexity measure as for the proteins (Methods)), even though every 16-mer has the same fitness.

The extraordinary strength of the bias towards high symmetry can be further illustrated by examining the prevalence of the two highest symmetry groups in the outcomes of evolutionary simulations. For 16-mers, there are 5 possible structures in class *D*_4_ (all symmetries of the square) and 12 in *C*_4_ (4-fold rotational symmetry). Even though these 17 structures represent just over a millionth of all 16-mer phenotypes, they make up about 30% of the structures that fix in the evolutionary runs, demonstrating an extraordinarily strong preference for high symmetry (See also Fig S3B). Comparing the histograms in Fig. 1C and Fig. 1F shows that the polyominoes exhibit a qualitatively similar bias towards high symmetry as seen for the proteins. We checked that this strong bias towards high symmetry/low 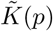 holds for a range of other evolutionary parameters (such as mutation rate) and for other polyomino sizes, see Fig S6 and SI section S3C. Natural selection explains why 16-mers are selected for (as opposed to other sizes). But, since every 16-mer is equally fit, natural selection does not explain the remarkable preference for symmetry observed here, which is instead caused by bias in the arrival of variation.

## Evolutionary simulations compared to sampling

In order to further understand the mechanisms that deliver the evolutionary preference for high symmetry, we calculated the probability *P*(*p*) of obtaining phenotype (polyomino shape) *p* by uniformly sampling 10^8^ genomes for the 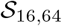 GP map, and counting each time a particular structure *p* (which can be any size) appears. Fig. 2 shows that *P*(*p*) varies over many orders of magnitude for different *p*. High *P*(*p*) only occurs for low 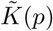 structures while high 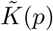 structures have low *P*(*p*). The inset of Fig. 2 shows that the *P*(*p*) from an evolutionary run from Fig. 1 closely follows the *P*(*p*) for 16-mers from random sampling. We tested this correlation for a range of different evolutionary parameters, and also for both randomly assigned and fixed fitness functions, and always observe relationships between *P*(*p*) and 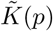 that are strikingly similar to those found for random sampling (Fig. S6).

**Fig. 2.**
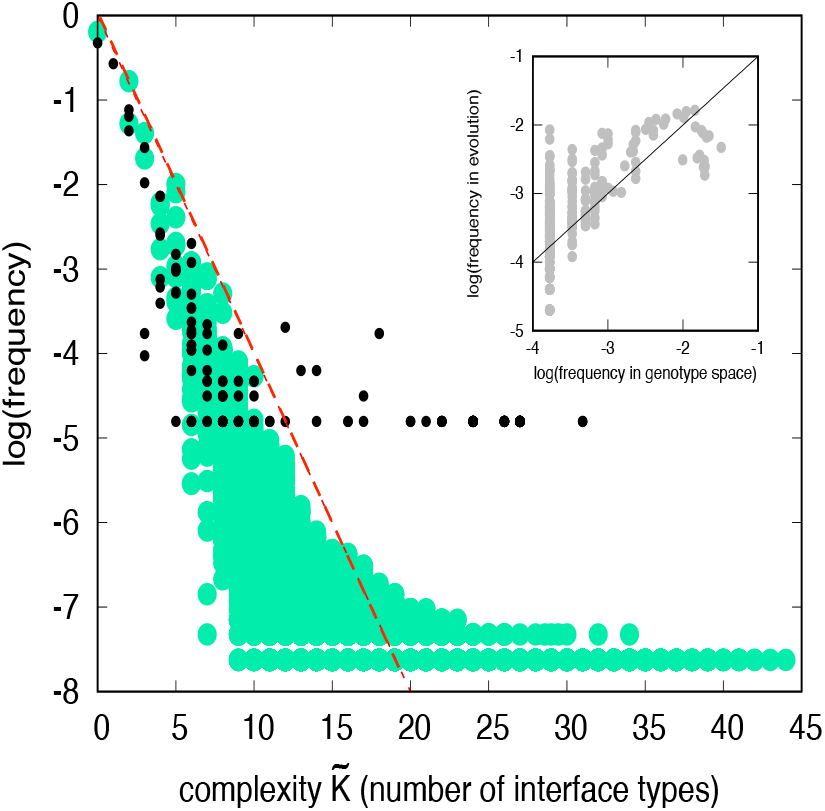
The frequency that a particular protein quaternary structure topology *p* (black circles) appears in the PDB versus complexity 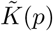=number of interface types, closely resembles the frequency distribution of all possible polyomino structures, obtained by randomly sampling 10^8^ genotypes for the 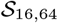 space (green cirles). Simpler (more compressible) phenotypes are much more likely to occur. An illustrative AIT upper bound from Eq. (1) is shown with *a* = 0.75, *b* = 0 (dashed red line). *Inset*. The frequency with which particular 16-mers are found to fix in evolutionary runs from Fig. 1E is predicted by the frequency with which they arise on random sampling of genotypes; the solid line denotes *x* = *y*.

The observed similarity in all these different evolutionary regimes is predicted by the *arrival of the frequent* population dynamics framework of ref [12] (SI section S2). For highly biased GP maps, it predicts that, for a wide range of mutation rates and population sizes, the rate at which variation (phenotype *p*) arises in an evolving population is, to first order, directly proportional to the probability *P*(*p*) of it appearing upon uniform random sampling over genotypes. Strong bias in the arrival of variation can overcome fitness differences, and so control evolutionary outcomes [12, 19]. Interestingly, recent results for deep learning support this evolution dynamics picture. Deep neural nets show a strong Occam’s razor like bias towards simple outputs [20] upon random sampling of parameters, and these frequent (and simple) outputs appear with similar probability under training with stochastic gradient descent [21]. This similarity between random sampling and the outcome of a stochastic optimiser strengthens the case for extending the applicability of the arrival of the frequent framework for highly biased to maps to a wide range of fitness landscapes (see SI section S2 for fuller discussion).

Fig. 2 also illustrates a striking similarity between the probability/complexity scaling for polyominoes and that of protein complex structures. Note that finite sampling effects lead to a widening of the lowest frequency outputs [8] (see also Fig. S5) suggesting that as more structures are deposited in the 3DComplex database [14] the agreement with the polyomino distribution may improve further. Given the simplicity of the polyomino model, this near quantitative agreement is probably somewhat fortuitous. Nevertheless, the arrival of the frequent mechanism, which for polyominoes explains the remarkably close similarity of the *P*(*p*) v.s. 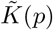 relationships across different evolutionary scenarios (see e.g. Figs S4–S9). predicts that the probability-complexity relationships for the protein complexes will be robust, *on average*, to the many different evolutionary histories that generated these complexes. Taken together, the data and arguments above strongly favour our hypothesis that bias in the arrival of variation, and not some as yet undiscovered adaptive process, is the first order explanation of the prevalence of high symmetries in protein complexes.

## Algorithmic information theory and GP maps

These results beg another question: Is the bias towards simplicity (low 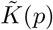) observed for protein clusters and polyominoes a more general property of GP maps? Some intuition can be gleaned from the famous trope of monkeys typing at random on typewriters. If each typewriter has *M* keys, then every output of length *N* has equal probability 1*/ M ^N^*. By contrast, if the monkeys’ keyboards are connected to a computer programming language then, for example, accidentally hitting the 21 characters of the program *print “01” 500 times;* will generate the *N* = 1000 digit string 010101 … with probability 1*/M* ^21^ instead of 1*/M* ^1000^. In other words, when searching in the space of algorithms, outputs that can be generated by short programs are exponentially more likely to be produced than outputs that can only generated by long programs.

This intuition that simpler outputs are more likely to appear upon random inputs into a computer programming language can be precisely quantified in the field of AIT [7], where the Kolmogorov complexity *K*(*p*) of a string *p* is formally defined as the shortest program that generates *p* on a suitably chosen universal Turing machine (UTM). While GP maps are typically not UTMs, and strictly speaking Kolmogorov complexity is uncomputable, a relationship between the probability *P*(*p*) and a computable descriptional complexity 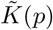 (typically based on compression) which approximates the true *K*(*p*) has recently been derived [8] for (non-UTM) input-output maps *f* : *I* → *O* between *N_I_* inputs and *N_O_* outputs. For a fairly general set of conditions, including that *N_I_N_O_*, and that the maps are asymptotically simple (see SI section S5), the probability *P*(*p*) that a map *f* generates output *p* upon random inputs can be bounded as:

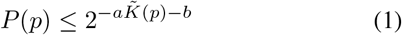

where 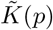 is an appropriate approximation to the true Kolmogorov complexity *K*(*p*), and *a* and *b* are constants that depend on the map, but not on *p*. While Eq. (1) is only an upper bound, it can be shown [22] that outputs generated by uniform random sampling of inputs are likely to be close to the bound. In extensive tests, Eq. (1) provided accurate bounds on the *P*(*p*) for systems ranging from from coupled differential equations to the RNA SS GP map [8] to deep neural networks [20], suggesting widespread applicability.

Since the number of genotypes is typically much greater than the number of phenotypes [1–3], and their relationship is encoded in a set of biophysical rules that typically depend weakly on system size, many GP maps satisfy the conditions [8] for Eq. (1) to apply (see also SI section S5). In Fig. 2, we we show an example of how Eq. (1) can act as an upper bound to *P*(*p*) for the polyominoes and the protein complexes. In SI section S5C, we demonstrate that this AIT formalism also works well for other choices of the complexity 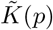, so that our results do not depend on the particular choices we make here. The AIT formalism also suggests that related systems should have similar probability-complexity relationships, which helps explain why the polyominoes and proteins have similar *P*(*p*) v.s. 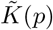 plots.

Since many GP maps satisfy the conditions for simplicity bias, including those where symmetry may be harder to define, we therefore hypothesised that a bias towards simplicity may also strongly affect evolutionary outcomes for many other GP maps in nature. We tested this hypothesis for RNA secondary structure and a model GRN.

## Simplicity bias in RNA secondary structure

Because it can fold into well-defined structures, RNA is a versatile molecule that performs many biologically functional roles besides encoding information. While sequence to 3D structure prediction is hard to solve computationally, a simpler problem of predicting secondary structure (SS), which describes the bonding pattern of the bases, can be both accurately and efficiently calculated [24]. The map from sequences to SS is perhaps the best-studied GP map, and has provided many conceptual insights into the role of structured variation in evolution [1–3, 12, 25–27]. It has already been shown, see e.g. [26–28], that the highly biased RNA GP map strongly determines the distributions of RNA shape properties in the fRNAdb database [23] of naturally occurring noncoding RNA (ncRNA). Although natural selection still plays a role (see [26, 27] for further discussions), the dominant determinant of these structural properties is strong bias in the arrival of variation [12]. It was recently shown [8] that the RNA SS GP map is well described by Eq. (1). Combining these observations leads to the hypothesis that functional ncRNA in nature should also be exponentially biased towards more compressible low 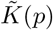 structures.

To test this hypothesis, we first, for length *L* = 30, calculate 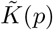 with a standard Lempel-Ziv compression technique [8] to directly measure the descriptional complexity of the dot-bracket notation of a SS (Methods and SI section S4). Fig. 3A shows that there is a strong inverse correlation between frequency and complexity for both naturally occurring and randomly sampled phenotypes (note that *L* = 30 is quite short so that finite size effects are expected [8] to affect the correlation with Eq. (1)). For longer RNA, the agreement with Eq. (1) is better (see e.g. ref [8], and Figs. S10, and S11. For *L* = 30 there are about 3×10^6^ possible SS [26], but only 17,603 are found in the fRNAdb database [23], and these are much more likely to be more compressible low 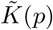 structures. Fig. 3C shows that randomly sampling of sequences provides a good predictor for the frequency with which these structures are found in the database, consistent with previous observations [26, 27] and the arrival of the frequent frame-work [12].

**FIG. 3.**
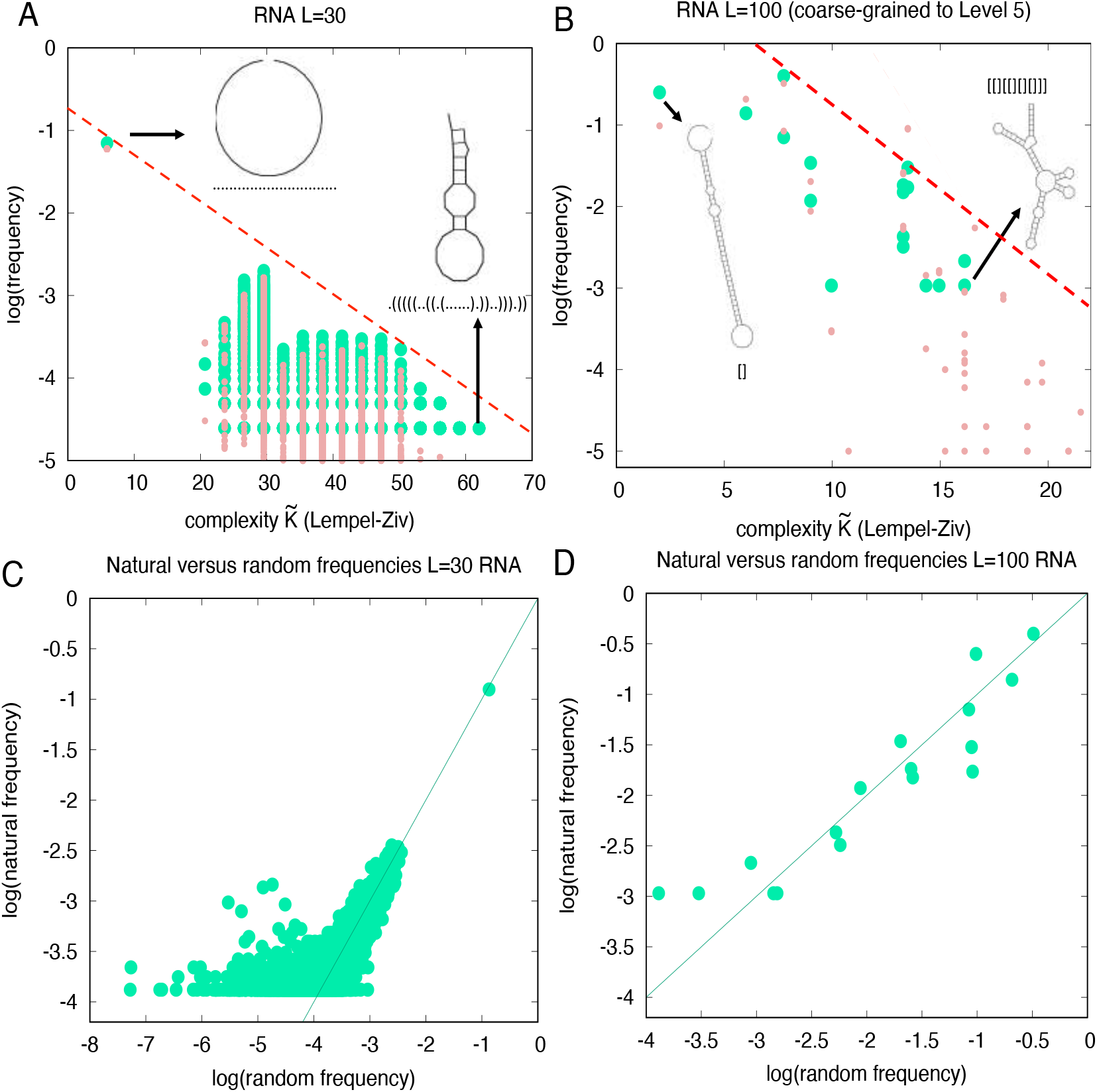
Frequency/probability versus complexity 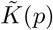 for (A) *L* = 30 RNA full SS and (B) *L* = 100 SS coarse-grained to level 5 (Methods). Probabilities for structures taken from random sampling of sequences (light red) compare well to the frequency found in the fRNA database [23] (green dots) for 40,554 functional *L* = 30 RNA sequences with 17,603 unique dot-bracket SS and for 932 natural *L* = 100 RNA sequences mapping to 16 unique coarse-grained level 5 structures. The dashed lines show a possible upper bound from Eq. (1). Examples of high probability/low complexity and low probability/high complexity SS are also shown. In (C) and (D) we directly compare the frequency of RNA structures in the fRNAdb database to the frequency of structures upon uniform random sampling of genotypes for *L* = 30 SS and *L* = 100 coarse-grained structures respectively. The lines are *y* = *x*. Correlation coefficients are 0.71 and 0.92, for *L*=30 and *L*=100 respectively, with *p*-value< 10^−6^ for both. Sampling errors are larger at low frequencies.

For lengths longer than *L* = 30, the databases of natural RNAs show little to no repeated SS, so individual frequencies cannot be extracted. To make progress, we apply a well established coarse-graining strategy that recursively groups together RNA structures by basic properties of their shapes [29], which was applied to naturally occurring RNA SS in ref. [27].

At the highest level of coarse-graining (level 5) there are many repeat structures in the fRNAdb database, allowing for frequencies to be directly measured (Methods). For *L* = 100 we compare the empirical frequencies to *P*(*p*) estimated by random sampling. Fig. 3B shows that there is again a strong negative correlation between frequency and complexity. (see also SI Tables II and III and Figs. S10 & S11 for fRNAdb and Rfam database data). Fig. 3D shows that natural frequencies are well predicted by the random sampling, as seen in ref. [27] for other lengths. Again, only a tiny fraction (≈ 1/10^8^) of all possible phenotypes is explored by nature [27]. The RNA SS GP map exhibits simplicity bias phenomenology similar to the protein complexes and the polyomino GP map. While the simpler group-theory based symmetries discussed for protein complexes and polyominoes do not apply here, the bias towards lower 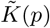 reflects the more generalised symmetries in the RNA SS structures.

## Model gene regulatory network

The protein and RNA phenotypes both describe shapes. Can a similar strong preference for simplicity be found for other classes of phenotypes? To answer this question, we also studied a celebrated model for the budding yeast cell-cycle [30], where the interactions between the biomolecules that regulate the cell-cycle are modelled by 60 coupled ordinary differential equations (ODEs) As a proxy for the genotypes, we randomly sample the 156 biochemical parameters of the ODEs (Methods). For each set of parameters, we calculate the complexity of the concentration versus time curve of the CLB2/SIC1 complex (a key part of the cycle) using the up-down method [31]. Fig. 4 shows that *P*(*p*) exhibits an exponential bias towards low complexity time curves, as hypothesised. Of course many of these phenotypes may not supply the biological function needed for the budding yeast cell-cycle. But interestingly, the wild-type phenotype has the lowest complexity of all the phenotypes we found, and is also the most likely to arise by random mutations. While the evolutionary origins of this GRN are complex, we again suggest that a bias towards simplicity in the arrival of variation played a key role in its emergence.

**FIG. 4.**
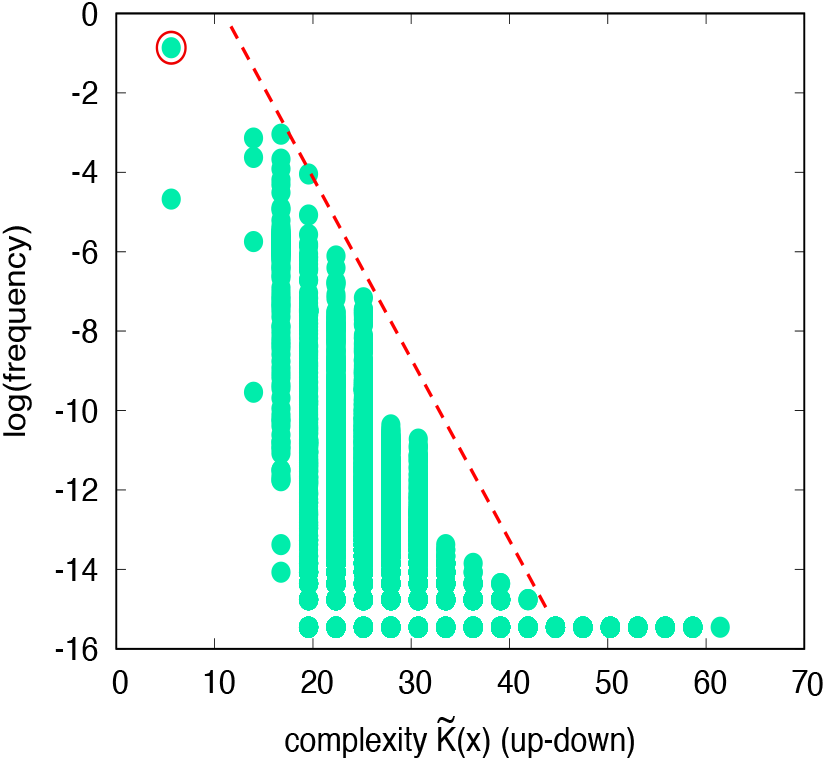
*P*(*p*) v.s. complexity 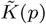 for the budding yeast ODE cell cycle model [30]. Phenotypes are grouped by complexity of the time-output of the key CLB2/SIC1 complex concentration. Higher *P*(*p*) means a larger fraction of parameters generate this time-curve. The red circle denotes the wild-type phenotype, which is one of the simplest and most likely phenotypes to appear. The dashed show a possible upper bound from Eq. (1). There is a clear bias towards low complexity outputs.

## Discussion

Our two main hypotheses are: 1.) GP maps are, under random mutations, exponentially biased towards phenotypic variation with low descriptional complexity, as predicted by AIT [8]. 2.) Such strong bias in the arrival of variation can affect adaptive evolutionary dynamics, leading to a much higher prevalence of low complexity (high symmetry) phenotypes than can be explained by natural selection alone.

The arguments above are general enough to suggest that many biological systems, beyond the examples we provided, may favour simplicity and, where relevant, high symmetry, without requiring selective advantages for these features. For example, there are claims that HP lattice proteins with larger *P*(*p*) are typically more symmetric [32], and similar patterns have been suggested for protein tertiary structure in the PDB [33]. In SI section S6 we present further evidence that protein tertiary structure, signalling networks [34] and Boolean threshold models for GRNs [35] also exhibit bias in the arrival of variation. At a more macroscopic level, a model of tooth development [36] suggests that simpler phenotypes evolved earlier, consistent with a high encounter probability in evolutionary search. Similarly, for both teeth [37] and leaf shape [38], mutations to more simpler tooth phenotypes are more likely than mutations to more complex phenotypes, an effect our theory also predicts. A recent theoretical study [39] of the development of morphology, which also found that simple morphologies were more likely to appear than complex ones upon random parameter choices. The L-systems used to model plant development [4] show simplity bias [8], and Azevedo *et al.* [40] showed that developmental pathways for cell lineages are significantly simpler (in a Kolmogorov complexity sense) than would be expected by chance.

On the other hand, for complex phenotypic traits affected by many loci, variation may be more isotropic so that bias is weak. For such traits, where classical population genetics – which focuses on shifting allele frequencies in a gene pool where standing variation is abundant – typically works well, our arguments may no longer hold. The phenotype bias we discuss here is fundamentally about the origin of novel variation [19, 41], and so is most relevant on longer time-scales.

Finally, simple systems have a larger *P*(*p*) and are therefore more mutationally robust [1–3, 26, 42] (see also SI section S4B). A correlation between low complexity and robustness is also found in the engineering literature [42, 43]. Biological complexity often arises from connecting existing components together into modular wholes. If the individual components are more robust, then it is easier for them to evolve additional function, for example a patch to bind to another protein, without compromising their core function. Similarly, a larger robustness may also enhance the ability of a system to encode cryptic variation, facilitating access to new phenotypes [44]. A natural tendency towards simpler and more robust structures may therefore facilitate the emergence of modularity, where individual components can evolve independently [45], and so make living systems more globally evolvable.

## METHODS

### Protein-complex topologies

Our analysis of protein quaternary structure builds upon the techniques and data presented in ref. [16], where a curated set of 30,469 monomers, 28,860 homomers, and 5,527 heteromers were extracted from the Protein Data Bank (PDB), and classified into 120 distinct topologies. These were then used to make a periodic table of possible topologies. Protein complexes are described in terms of a weighted subunit interaction graph. An illustration of the topologies, and how they are generated is shown in Fig S1, for two heteromeric complexes, and their final graph topologies. Further examples of topologies and the PDB structures they describe can be found at http://www.periodicproteincomplexes.org/. The nodes of the graph are labelled according to their protein identities and the weights of the connections are the interface sizes in Å^2^. The procedure for enumerating possible topologies, and for classifying existing and potential topologies is described more fully in ref. [16]. This approach only considers the largest interfaces, which if cut would disconnect the complex. The reason is that small interfaces that can be cut without disconnecting the complex are likely to be circumstantial, and unlikely to play an important role in the assembly and evolution of the complex. After constructing the weighted subunit interaction graphs in this manner we identify the topologically distinct interaction graph of subunit types (see for example Fig. S1C, with the additional distinction between symmetric and asymmetric self-interactions of a subunit type, corresponding to homomeric interfaces.

We take the number of interface types of protein complex *p* to be the complexity measure 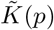. This choice is proportional to the number of individual mutations needed to generate the self-assembled complex. See SI section S5C for a longer discussion of different possible complexity measures. Unlike the polyomino case, where the building block is a square tile, the geometry of an individual protein is highly variable. For example, a cyclic homomeric 6-ring and a cyclic homomeric 10-ring will have the same topologically distinct interface configuration ( which is just the two parts of the same asymmetric interface on a single subunit). This will be distinct from a heteromeric 6-ring in which we have two halves of two different symmetric interfaces on a subunit, and also distinct from a simple heterodimer. All three of these however have the same number of interface types (2) and so appear at 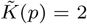 in the distribution of Figs. 1 and 2 in the main text. The single point that appears at 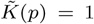 for Fig. 2 is a homodimer and the single point at 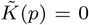 is a monomer. The symmetries of all protein complexes presented here are taken directly from the PDB.

To calculate the symmetries of all hypothetical protein complexes of size six in Fig. 1C, we used the following procedure. We first consider all topologically distinct graphs of size six with up to six different subunit types and symmetric or asymmetric homomeric interfaces between subunits of the same type. By comparing all 6! possible permutations of the adjacency matrix and the associated node labels we then calculate the permutation symmetries of the node types on these graphs as a proxy for the spatial symmetry of the hypothetical protein complexes that they represent. This collapses D3 and C6 into one category, but allows us to distinguish this category from C3, C2, and C1 (and these from each other). Further discussion of the protein complexes can be found in SI section S1.

### Polyominoes

The polyomino model was implemented as described in refs. [6, 17, 18]. The genome encodes a ruleset consisting of 4*n* numbers which describe the interactions on each edge of *n* square tiles. Each number is represented as a length *b* binary string, so that the whole genome is a binary string of length *L* = 4*nb*. The interactions bond irreversibly and with equal strength in unique pairs (1 ↔ 2, 3 ↔ 4, …), with types 0 and 2*^b^* − 1 being neutral, not bonding to any other types. We label a given polyomino GP map with up to *n* possible tiles and 4*n* possible colors as 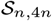 ; in this paper we usually work with 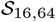

The assembly process is initiated by placing a single copy of the first-encoded subunit tile on an infinite g rid. A different protocol where any tile may be used to seed the assembly is also possible and does not significantly affect the results presented here. Assembly then proceeds as follows. 1) Available moves are identified, consisting of an empty grid site, a particular tile and a particular orientation, such that placing that tile in that orientation in the site will form a bond to an adjacent tile that has already been placed. If there are no available moves, terminate assembly. 3) Choose a random available move and place the given tile in that orientation at that site. 4) If the current structure has exceeded a given cutoff size, terminate assembly. 5) Go to step 1.

This process is repeated 20 times to ensure that assembly is *deterministic* – that is, that the same structure is produced each time. If different structures are produced, or the structure exceeds a cutoff size (here taken to be larger than a 16×16 grid), the structure is placed in the category ‘UND’ (unbounded or non-deterministic). For the calculation of probabilities/frequencies *P*(*p*) we ignore genotypes that produce the UND phenotype. This choice mimics the intuition that unbounded protein assemblies, or else proteins that do not robustly self-assemble into the same shape, are highly deleterious.

The ruleset 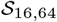 allows any 16-mer to be made, since it is always possible to use addressable assembly where each tile is unique to a specific location. But many 16-mers can be made with significantly fewer than 16 tile types, although there are examples that (to our knowledge) can only be made with all 16 tiles, so that a space allowing up to 16 tiles is needed.

To assign complexity values for the polyominoes, a measure similar to that used for the proteins was applied. First, the minimal complexity over the different genomes that generate polyomino *p* is estimated by a sampling and finding the shortest rule set and removing redundant information. The search for a minimal complexity genome will be more accurate for high probability polyominoes than for low probability polyominoes. We checked that for most structures only a fairly limited amount of sampling provided an accurate estimate of the minimal complexity; the minimal complexity genome is typically the most likely to be found. The effects of finite sampling are illustrated further in SI section S3 and Fig. S5. The complexity 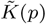 of polyomino *p* is then given by the smallest number of unique edge labels (interface types) in the minimal genomes – thus, twice the number of hetero-interactions, just as in the protein system above. A longer discussion of different choices of complexity measure, showing that the qualitative behaviour is not very sensitive to details in the choice of approximate measure of algorithmic (Kolmogorov) information, can be found in SI Section S3C.

Evolutionary simulations of polyomino structures are performed following methods described in ref. [17]: A population of *N* binary polyomino genomes is maintained at each time-step. The assembly process is performed for each genome and the resulting structure is recorded. UND genomes are assigned zero fitness. Other structures are assigned a fitness value based on the applied fitness function. These fitness values are used to perform roulette wheel selection, whereby a genome *g_i_* with fitness *f* (*g_i_*) is selected with probability *f* (*g_i_*)/ Σ_*j*_ *f* (*g_j_*). Selection is performed *N* times (with replacement) to build the population for the next time-step. Selected genomes are cloned to the next generation, then point mutations are applied with probability *μ* at each locus. A point mutation changes a 0 to a 1 and vice versa in the genome. We do not employ crossover or elitism in these simulations.

We employ several different fitness functions. In the *unit fitness* protocol, all polyomino structures that are not UND are assigned fitness 1. In the *random fitness* protocol, each polyomino structure is assigned a fitness value uniformly randomly distributed on [0, 1], and these values are reassigned for each individual evolutionary run. In the *size fitness* protocol, a polyomino of size *s* has fitness 1*/*( |*s − s*\*| + 1), so that polyominoes of size *s*\* have unit fitness and other sizes have fitness decreasing with distance from *s*\*. The simulations for Fig. 1E were done with *N* = 100 and *μ* = 0.1 per genome, per generation. A number of other evolutionary parameters are compared in SI section 3C and Fig. S6, showing that our main result – that the outcome of evolutionary dynamics exhibit an exponential bias towards simple structures – is not very sensitive to details such as mutation rate or the choice of fitness function.

### RNA secondary structure GP map

For *L* = 30 RNA we randomly generated 32,000 sequences, and for *L* = 100, we generated 100,000 random sequences. As in refs. [26, 27], secondary structure (SS) is computationally predicted using the `fold` routine of the Vienna package [24] based on standard thermodynamics of folding. All folding was performed with parameters set to their default values (in particular, the temperature is set at *T* = 37°C). We then calculated the neutral set size (NSS(p)), the number of sequences mapping to a SS *p*, for each SS found by random sampling, by using the neutral network size estimator (NNSE) described in ref [28], which is known to be quite accurate for larger NSS structures [26]. We used default settings except for the total number of measurements (set with the -m option) which we set to 1 instead of the default 10, for the sake of speed, but this does not noticeably affect the outcomes we present here.

RNA structures can be represented in standard dot-bracket notation, where brackets denote bonds, and dots denote unbonded pairs. For example, …((…))…… means that the first three bases are not bonded, the fourth and fifth are bonded, the sixth through ninth are unboded, the tenth base is bonded to the fifth base, the eleventh base is bonded to the fourth base, and the final four bases are unbounded. For shorter strands such as *L* = 30, the same SS can be found multiple times in the fRNAdb.

For longer strands, finding multiple examples of the same SS becomes more rare, so that SS frequencies cannot be easily directly extracted from the fRNAdb. However, it seems reasonable, especially for larger structures, that fine details of the structures are not as important as certain more gross structural features that are captured by a more coarse grained picture of the structure. In this spirit, we make use of the well known RNA abstract shape method [29] where the dot-bracket SS are abstracted to one of five hierarchical levels, of increasing abstraction, by ignoring details such as the length of loops, but including broad shape features. For the *L* = 100 data we choose the fifth, or highest level of abstraction which only measures the stem arrangement. This choice of level is needed to achieve multiple examples of the same structure in the fRNAdb database, so that a frequency can be directly determined with statistical significance. The SS were converted to abstract shapes with the online tool available at https://bibiserv.cebitec.uni-bielefeld.de/rnashapes. Using these coarse-grained structures means that the theoretical probability *P*(*p*) can be directly calculated from random sampling of sequences, where *N_G_* is the number of sequences, which for an RNA GP map for length *L* RNA is given by *N_G_* = 4*^L^*. A similar calculation of the *P*(*p*) for RNA structures for *L* from 40 to 126 at different levels of coarse-graining can be found in [27].

To generate the distributions of natural RNA we took all available sequences of *L* = 30 and *L =*100 from the non-coding functional RNA database (fRNAdb [23]). As in ref. [26], we removed a small fraction (~ 1%) of the natural RNA sequences containing non-standard nucleotide letters, e.g. ‘N’ or ‘R’ because the standard folding packages cannot treat them. Similarly, a small fraction (~ 2%) of sequences were also discarded due to the neutral set size estimator (NSSE) failing to calculate the NSS (this is only relevant for *L* = 30). We have further checked that removing by hand any sequences that were assigned putative roles, or are clear repeats, does not significantly affect the strong correlation between the frequencies found in the fRNA database and those obtained upon random sampling of genotypes. For a further discussion of the question of how well frequency in the databases tracks the frequency in nature, see also refs. [26, 27] and Fig. S10 where a comparison with the Rfam database is also made. Note that the similar behaviour we find across structure prediction methods, strand lengths, and databases would be extremely odd if artificial biases were strong on average in the fRNA database. We used 40, 554 unique RNA sequences of *L* = 30, taken from the fRNAdb, corresponding to 17,603 unique dot-bracket structures. Similarly, we used 932 unique fRNAdb *L* = 100 RNA sequences, corresponding to 17 unique level 5 abstract structures/shapes.

To estimate the complexity of an RNA SS, we first converted the dot-bracket representation of the structure into a binary string *p*, and then used the Lempel-Ziv based complexity measure from ref. [8] to estimate its complexity. To convert to binary strings, we replaced each dot with the bits 00, each left-bracket with the bits 10, and each right-bracket with 01. Thus an RNA SS of length *n* becomes a bit-string of length 2*n*. Because level 5 abstraction only contains left and right brackets, i.e. [ and ], we simply convert left-bracket to 0, and right to 1 before estimating the complexity of the resulting bit string via the Lempel-Ziv based complexity measure from ref. [8]. The level 5 abstract trivial shape with no bonds is written as underscore, and this we simply represented as a single 0 bit. SI section S4 provides more background on RNA structures, and section S5B more detail of the complexity measure.

### GRN of budding yeast cell-cycle

The budding yeast (*S. cerevisiae*) cell-cycle GRN system from ref. [30] consists of 60 coupled ordinary differential equations (ODEs) relating 156 biochemical parameters. The model parameter space (i.e. genotype space) was sampled by picking random values for each of the parameters by multiplying the wild-type value by one of 0.25, 0.50, …, 1.75, 2.00, chosen with uniform probability. The ODEs generate concentration-time curves for different biochemicals involved in cell-cycle regulations. All runs were first simulated for 1000 time steps, with every time step corresponding to 1 minute. Next, we identified the period of every run (usually on the order of 90 time steps), took one full oscillation and coarse-grained it to 50 time steps. This way, if two genotypes produce curves which are identical up to changes in period, they should ultimately produce identical or nearly-identical time series and binary string phenotypes. For every “genotype” or set of parameters, the curves for the CLB2/SIC1 complex are then discretised into binary strings using the “up-down” method [31]: for every discrete value of *t = δt,* 2*δt,* 3*δt, …*, we calculate the slope *dy/dt* of the concentration curve, and if *dy/dt* ≥ 0, a 1 gets assigned to the *j*-th bit of the output string, otherwise, a 0 is assigned to it. All strings with the same up/down profile were classified as one phenotype. To generate the *P*(*p*) in Fig. 4, 5 × 10^6^ inputs were sampled. Complexity 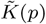 is assigned by using the Lempel Ziv measure from ref. [8] (see also SI section S5B) applied to binary output strings. As shown in ref. [8], this methodology works well for coupled differential equations, and the choice of input discretisation, sample size and initial conditions does not qualitatively affect the probability-complexity relationships obtained. The wildtype curve can be observed in in Fig. 2 of ref. [30] where it is labelled Clb2*_T_*.

## Author Contributions

IGJ and SG did the polyomino simulations, SEA generated the protein data, KD generated the RNA data, KD and CQC analysed the GRN, JPKD, SEA and AAL supervised the polyomino work, AAL supervised the RNA and GRN work. All authors helped analyse the data and write the paper.

## Acknowledgements

We thank N. Martin, V. Mohanty, J. Bohlin, S. Schaper and H. Zenil for discussions.

## S1. SUPPLEMENTARY TEXT FOR PROTEIN QUATERNARY STRUCTURE

While the tertiary structure of a protein describes the folded state of an individual polypeptide chain, the quaternary structure of a protein describes how individual protein subunits bond toform the final c omplex. Over half the proteins found in nature (sofar) form homomeric or heteromeric complexes with other proteins, and these structures tend to be highly conserved on evolutionary time-scales [2, 3]. Interestingly, the physical assembly pathways may mimic the evolutionary pathways that led to a particular protein assembly [4, 5]. In ref. [1] the authors combined bioinformatic searches together with electrosplay mass spectrometry experiments to hypothesise that protein complex topologies can be generated from combinations of three basic types of assembly steps, namely dimerisation, cyclisation, and heteromeric sub-unit addition. These combinations were used to generate a periodic table of (theoretically) possible topologies. Most protein complex topologies found in the PDB were shown to fall into one of the topologies predicted from their procedure, and thus can be classified into their periodic table. At the same time, many potential topologies have not (yet) been found, and most of these are complex, that is they would need a larger complexity 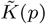 to be produced than the values we measure for existing structures. This fact is consistent with our hypothesis that complex topologies are harder to evolve than simpler ones are, and so less likely to appear in nature.

In Fig. S1, we illustrate how the methods from ref. [1] are used to generate topologies and complexities of the proteins structures.

A key question is whether or not the patterns we observed in the PDB mirror patterns seen more widely in the biological world. While this question is hard to answer *a-priori*, we note that it would be extremely surprising if the strong bias towards symmetry were merely an artefact of the PDB. To further test this question, we follow methodology from ref. [1] by applying a stringent redundancy filter that removes multiple copies of the same protein from the data if they share more than 50% sequence identity. This filter is used for a subset of the analysis conducted in [1], and the filtered subset of complexes is provided as part of the Supplementary Information of that publication. As can be seen when comparing Fig. S2 (A) to Fig. 1B in the main text, this procedure reduces the amount of data significantly from 1778 to 545 6-mer topologies, but the overall trend is very similar, supporting our hypothesis that the global bias towards symmetry observed in the PDB mirrors a broader trend in biology, and is not a database artefact.

**FIG. S1.**
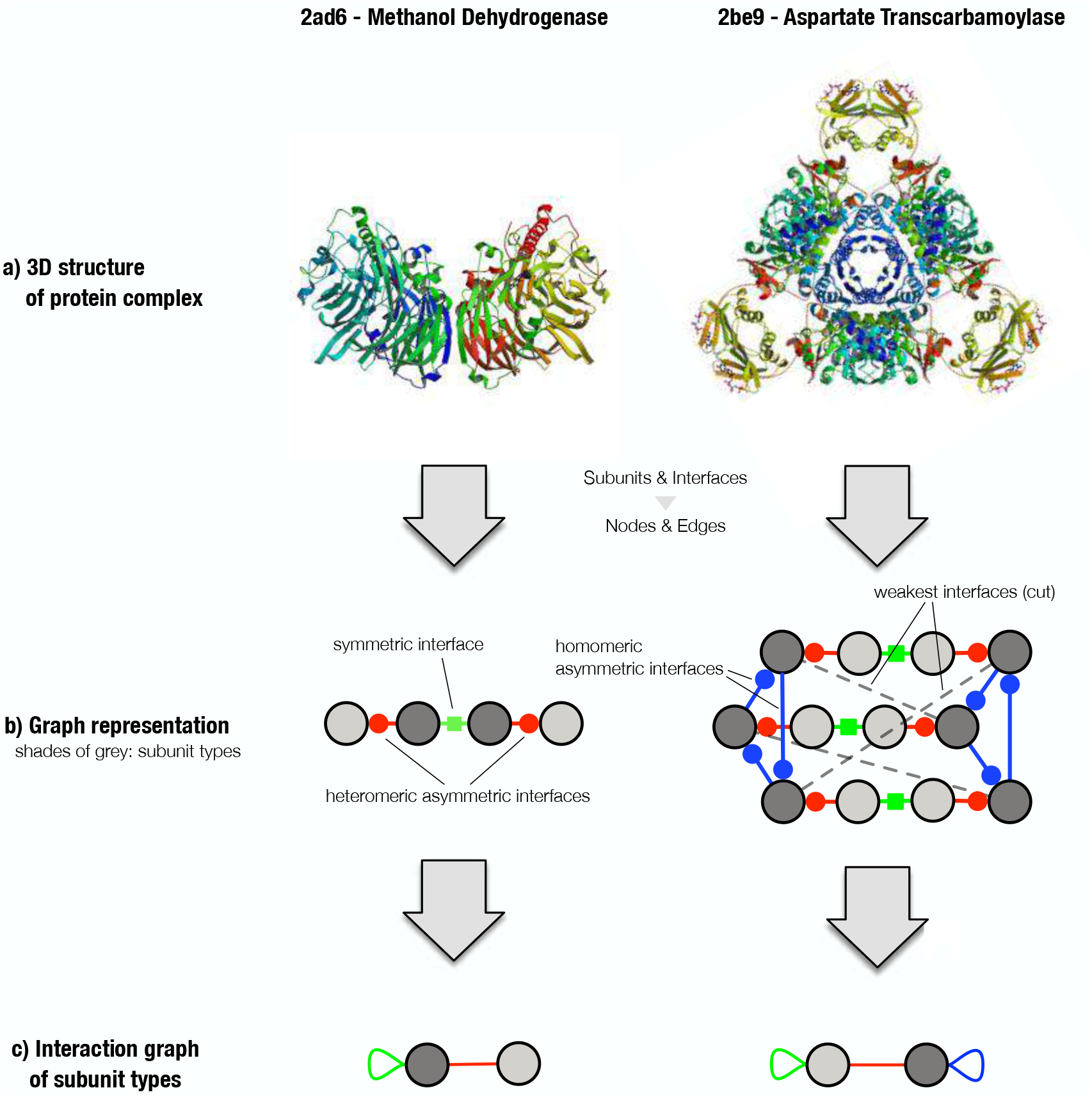
Graph representation of protein-complex topologies following the methods from ref. [1]. (a) Two heteromeric complexes, a 4-mer methanol dehydrogenase (2ad6), and a 12-mer aspartate transcarbamoyfase (2b39). (b) Symbolic graph representations of the proteins and the interfaces between them. Different shades of grey denote different types of protein subunit (according to sequence similarity). Interface colours denote symmetric (green), homomeric asymmetric (blue), and heteromeric asymmetric (red) interlaces. Weak interfaces that are disregarded (see Methods section IA) are shown as dashed lines. (c) Interaction graph of subunit types, which is a reduced representation of the same complex that shows each type of distinct interface that exists in the complex exactly once. For instance, the green self-loop on the light grey node in the left column indicates the symmetric (green) interaction with another copy of the same subunit type in b), whereas the red connection denotes the heteromeric interaction between the two different subunit types. The complexity of a protein complex is given by the number of interface types, where symmetric interfaces (green) consist of one self-interacting type, and asymmetric interfaces (blue and red) consist of two types. The complexities of the examples shown are therefore 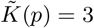 for 2ad6 and 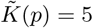 for 2b39.

**FIG. S2.**
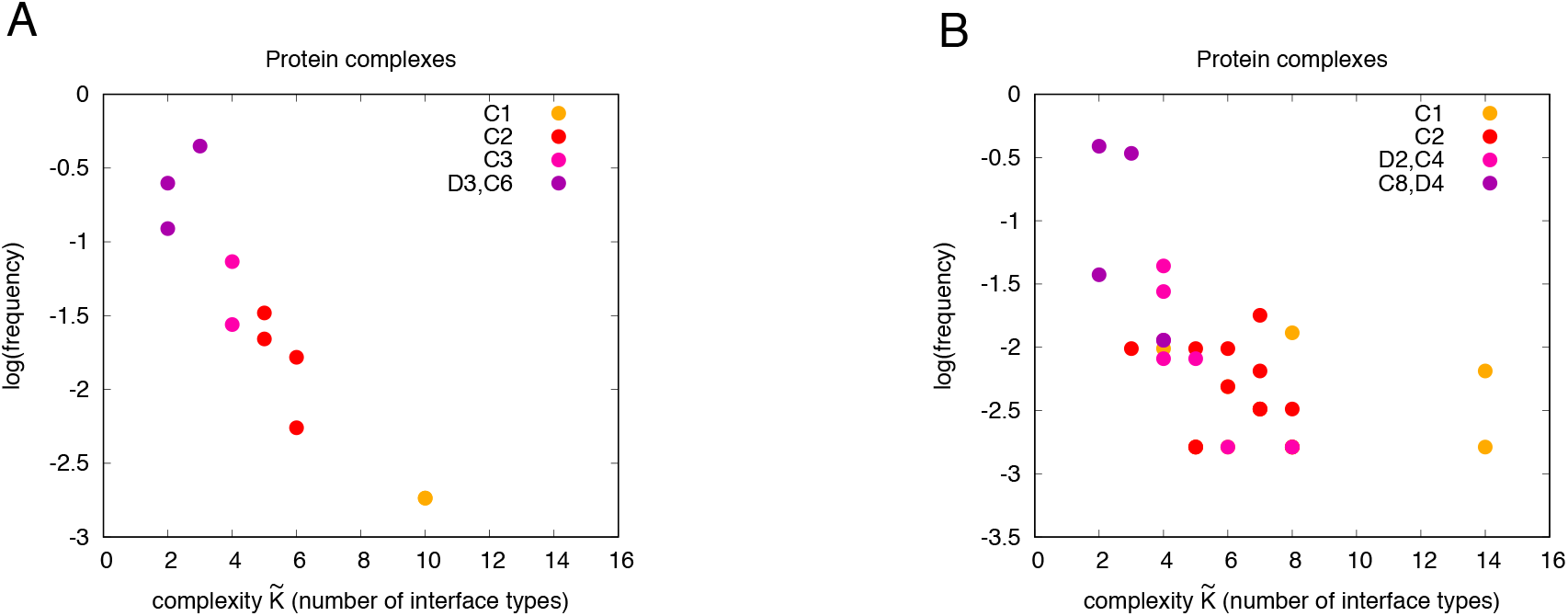
**(A)** Frequency of 6-mer protein complex topologies found in the PDB versus the number of interface types (a measure of complexity), as in Fig. 1B of the main text, but with a redundancy filter from [1] applied. **(B)** Frequency of 8-mer protein complex topologies versus complexity (no redundancy filter). For both sizes there is a strong preference for low complexity/high symmetry structures. Symmetries are in standard Schoenflies notation.

**FIG. S3.**
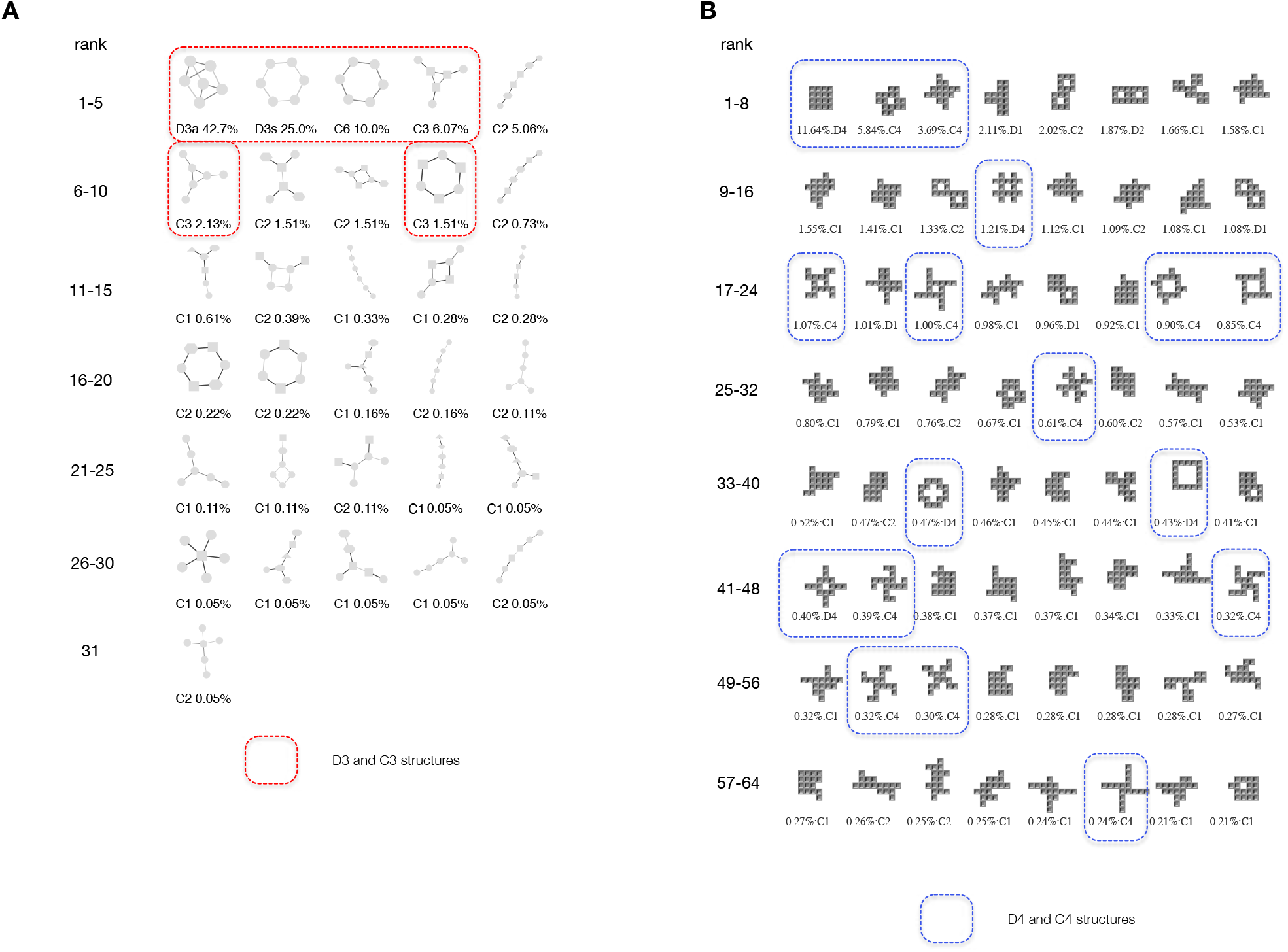
A) Protein 6-mer complex topologies from Fig 1B, ordered by their frequency in the PDB database. The highlighted ones are those with symmetry *C*_3_ or higher. The higher symmetry ones are significantly more likely on average to appear in the database than lower symmetry structures. Similar results are found for other sizes. Darker and paler lines corresponds to asymmetric and symmetric interfaces, respectively. **B) Most frequently found 16-mer polyomino structures from Fig 1E**. These 64 most likely structures (out of 13,079,234 possible 16-mers) include all 5 of the most symmetric polyominoes with *D*_4_ symmetry, and all 12 with the next symmetry (*C*_4_). Together these 64 structures make up about 64% of the total probability weight for this evolutionary run even though they only make up a tiny fraction ≈1/200, 000 of all possible structures. By contrast, while the low symmetry *C*_1_ structures make up 99.9% of all possible polyomino shapes, they are heavily suppressed in the evolutionary run relative to higher symmetry shapes. All 16-mer structures share the same fitness, so while the emergence of a 16-mer is due to natural selection, the strong bias towards simpler/more symmetric structures illustrated here is not caused by natural selection, but rather by the very large differences in rates of the arrival of variation.

The main text only shows data for 6-mers. We found similar trends for other sizes. As an example, in Fig, S2 (B) we show a frequency vs. complexity plot for 8-mers, which shows a similar frequency-complexity behaviour to that found for 6mers, and for other sizes as well.

In the main text, Fig. 1C, we showed that structures found in the PDB were highly biased towards a small subset of highly symmetric structures. In Fig. S3 (A) we show the 6-mer topologies in more detail, including their symmetries. This plot helps illustrate the main point observed in Fig. 1B, which is that highly symmetric structures are much more likely to appear in nature than structures with lower symmetry.

## S2. POPULATION DYNAMICS AND ARRIVAL OF THE FREQUENT FOR A HIGHLY BIASED GP MAP

Fig. 1 from the main text shows the results of an evolutionary simulation for polyominoes with a fitness function that has a maximum for size *s*\* = 16. We argue that, on average, the frequencies with which phenotype occur should be qualitatively similar to the probabilities *P*(*p*) that they obtain upon random sampling of sequences for a wide range of population dynamic parameters by invoking the “arrival of the frequent” population dynamics framework from ref. [6]. To those steeped in traditional population genetics, where bias in the introduction of variation is rarely taken into account [7], it may seem highly surprising that random sampling over genotypes should be able to make such predictions when a GP map is highly biased. We therefore briefly sketch out some key arguments from the literature from that framework below.

The first point to note is that given a genotype that maps to source phenotype *y*, the *mean* probability *ϕ_py_* that novel phenotype *p* ≠ *y* arises by a random point mutation can, to first order, be approximated quite well by the *global* probability *P*(*p*) that *p* appears upon random sampling of genotypes, regardless of the source phenotype *y*, an effect which has been observed for a range of GP maps [6, 8–10]. This average equivalence of the local probability that a phenotype emerges in a population upon mutations, with the global probability *P*(*p*) that it appears upon random sampling of sequences is what one might expect as a first order null model. While there will obviously be deviations from this simple rule, when the frequencies vary by many orders of magnitude, as they do for the GP maps studied here, and as the AIT formalism predicts will be true much more widely [11], then these deviations are likely to be relatively much smaller, on average than the effect of global frequency. If one is interested in global predictions, then the global frequency will be a good first order guide. This argument is crucial for understanding why we can take such a simple step as random sampling of sequences, and yet make predictions, on average, for outcomes that have arisen from many complex evolutionary histories. We show explicitly in Fig. S6C that this prediction of local frequency being a good predictor of the rate at which a phenotype arises in a population works remarkably well for polyomino simulations. In [6] the same effect is demonstrated for simulations of the *L* = 20 RNA map.

Interestingly the mutational robustness *ρ_y_*, defined as probability that a point mutation generates the same phenotype as the source *y*, shows much less variation, and scales as *ρ_y_* ~ log(*P* (*y*)) for all the GP maps that we are aware of [6, 8, 9, 12–15]. Since *P*(*p*) can vary over many orders of magnitude, the probability that a particular novel phenotype arises by mutations can vary by many orders of magnitude, depending to first order on its *P* ( *p*). By contrast, the robustness *ρ* varies less dramatically for a typical GP map.

Next consider a population with a carrying capacity of *N* individuals, a mutation rate of *μ* per site and a genome length of *L*. Then as shown in ref. [6], under the assumptions above, the average over evolutionary runs of the median time 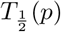 to discover a particular new phenotype *p* is is well described by the simple analytic form

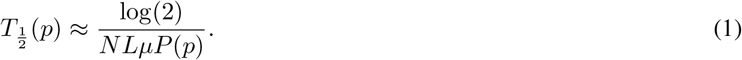

in polymorphic limit (*NLμ* ≫ 1), and by

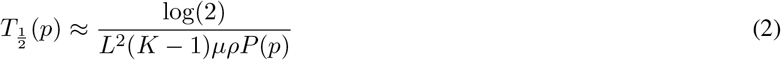

in the opposite monomorphic limit where *NLμ* ≪ 1. Here *K* is the alphabet size of the genome, and the robustness *ρ* is that of the neutral network the system is on. Extensive evolutionary simulations for a range of populations sizes and mutation rates showed that both these analytic forms quantitatively describe the discovery time 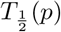 [6] in their respective regimes, and that one can smoothly interpolate between them for intermediate regimes.

The AIT upper bound on *P*(*p*), (Eq. (1) in the main text and Eq. (17) in this SM, allows us to derive lower bounds for the median discovery time in the polymorphic regime:

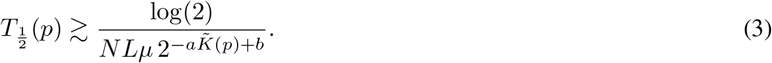

and in the monomorphic regime:

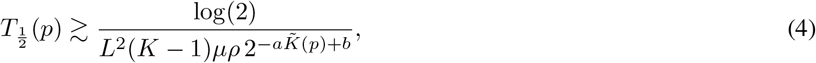

where we have ignored for now the scaling of *ρ* with the phenotype the system starts from, since this typically varies by less than an order of magnitude for phenotypes of interest, much less than the many orders of magnitude variation in *P*(*p*).

The observed variation in *P*(*p*) (or 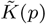) immediately implies that the discovery time 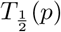 of a new phenotype *p* will typically vary over many orders of magnitude, with simple phenotypes being exponentially more likely to occur than more complex phenotypes. Since evolutionary time-scales for real populations are limited, typically only the most frequent phenotypes are likely to arise as potential variation for natural selection to act on. And thus it is only these frequent phenotypes that can fix, and that are therefore observed in nature. This *arrival of the frequent* effect [6] has already been observed for non-coding functional RNA where only a minescule fraction of all possible SS are observed in nature. Indeed those that are found can be shown to be frequent structures with relatively large NSS or *P*(*p*) [16, 17].

We note that this effect differs fundamentally from mechanisms such as the *survival of the flattest* [18], which implicitly assume that frequent and rare phenotypes are both present as potential variation. Instead, the arrival of the frequent is a non-ergodic and time-dependent effect that is fundamentally about the exponentially large range of rates at which novel variation arises. In the limit of infinite time all potential variation could appear, and then effects such as the survival of the flattest could be more relevant for these systems (see [6] for a discussion). However, that regime is not (even nearly) reached for the systems we study.

In sum, very strong biases in *P*(*p*) translate into equivalent biases in 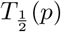, which, in turn, play an important role in evolutionary outcomes within this arrival of the frequent picture. When the range of *P*(*p*) is large, as we argue is generically the case for the GP maps of protein quaternary structure, RNA SS, and GRN that we study, then the arrival of the frequent effect may in fact dominate (on average) over various selective pressures.

See also refs. [7, 19] for a much broader discussion of the rate of introduction of new phenotypes in the evolutionary literature, and [20] for an important discussion of bias in the introduction of new variation.

In Fig. S4(a) we show some typical trajectories for a simulation with a fitness function that is maximal for 16-mers. In Fig. S4(b) we schematically illustrate the main idea of the arrival of the frequent [6].

In section S3C we illustrate with evolutionary simulations how the arrival of the frequent mechanism described above determines evolutionary outcomes for a range of population dynamic parameters, and different fitness functions. When this effect dominates, the outcomes only weakly depend on the details of the evolutionary dynamics. We recognize that this result may seem surprising to some, but the data we present is pretty unambiguous for these systems.

**FIG. S4.**
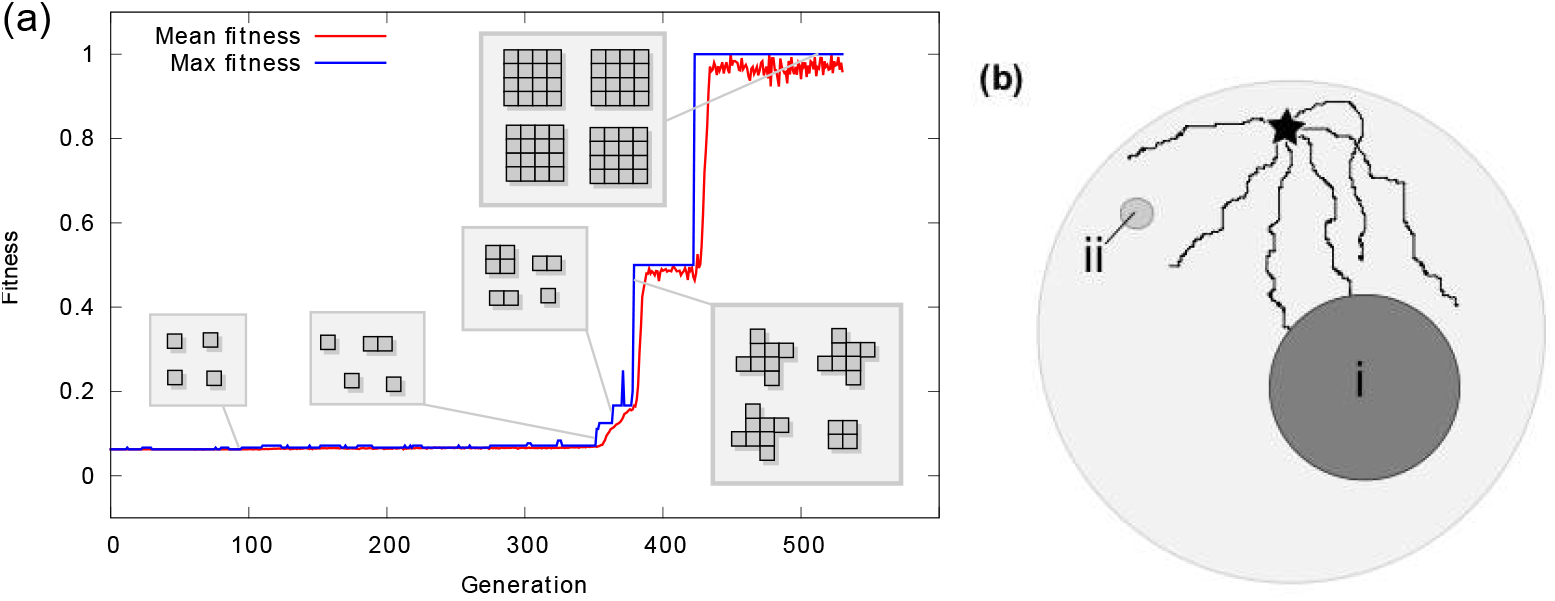
Schematic picture of evolutionary trajectories in search space. (a) An example of a typical evolutionary simulation run in the directed evolution model for *s*\* = 16 for *N* = 100 and *μ* = 0.1. Traces show mean and maximal fitness within a population, insets show a snapshot of a subset of the population at a given time. Starting from a population of simple structures, more complex structures evolve as evolutionary innovation discovers bonding approaches to produce fitter (larger) polyominoes. (b) A schematic figure illustrating how, if a population starts at phenotype (star), it is much more likely to find a phenotype such as **i** with a large NSS, than a phenotype such as **ii** which has a small NSS, even if the latter is more fit than the former. In the case of (a) and Fig. 1 in the main text, all 13,079,255 16-mers have the same maximum fitness, but a structure such as the square is much more likely to appear than a low symmetry structure because the square has a large NSS.

### A. Arrival of the frequent and the analogy between evolution on biased GP maps and deep learning

There is an interesting analogy between the arrival of the frequent in a highly biased evolutionary system, and training deep neural networks (DNNs) in the context of supervised learning [21]. Given a training set *S = {x_i_, y_i_*} of input-output pairs, where the *x_i_* are drawn from an input space 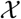, and the *y_i_* are drawn from an output space 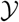. The supervised learning training task is to find a set of neural network parameters such that the DNN, upon the set of inputs {*x_i_} ∈ S*, produces a set of outputs 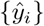 that minimises a loss function 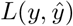. Zero training error means that 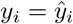 for all {*y_i_} ∈ S*. The most popular optimisation technique to find parameters that minimise the loss function is called stochastic gradient descent (SGD). This method follows the local gradient of the loss landscape. This problem is similar to an evolutionary system which maximise a local fitness function. For this reason, genetic algorithms, inspired by evolutionary dynamics, have been used to train DNNs, see e.g. [22] for a recent application for supervised learning, or [23] for an application of genetic algorithms to DNN based reinforcement learning.

DNNs can be viewed as function approximators, where the function *f* determines which outputs 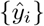 the DNN produces upon inputs {*x_i_*}. A parameter-function map can be defined as follows. The space of functions the model can express is a set 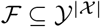. If the model takes parameters within a set *W* ⊆ ℝ^*n*^, then the parameter-function map 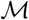 is defined as

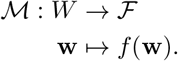

Since DNNs are typically employed in the overparameterized regime, with many more parameters than inputs or outputs, there are typically many different sets of parameters that map to the same function *f*. One can define a probability *P* (*f*) that a DNN produces a function *f* upon random sampling of its parameters (for simplicity, consider functions *f* restricted to the domain defined by the set of inputs *x_i_* of interest.). It has been recently shown [24] that, just as for GP maps, this parameter-to-function map follows the same AIT predicted simplicity bias scaling from Eq. (17). In other words, DNNs have a strong bias towards simple functions upon random sampling of parameters.

However, as mentioned above, the relevance of *P* (*f*) is not immediately obvious, since DNNs are not trained by random sampling of parameters, which would be very inefficient, but rather by SGD or related optimisation techniques which start with a particular set of parameters, and make local moves in parameter space to reach a loss minimum (equivalent to a fitness optimum for evolutionary systems). Interestingly, it has recently been shown for a wide range of data sets and for different loss functions, that training by SGD, which only takes into account local information on the loss function 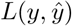, when averaged over initial conditions, generates functions *f* with a probability *P_SGD_*(*f*) ≈ *P* (*f*) [25]. This behaviour has as strong resemblance with what we find in this paper for evolutionary systems, where the global probability *P*(*p*) that a particular phenotype *p* appears upon random sampling is predictive of an evolutionary process that only locally samples a fitness function. Both the DNN parameter-function map, and the evolutionary GP maps, are highly biased. For such highly biased systems, the arrival of the frequent, or closely related phenomenology, explains why random sampling of parameters predicts outcomes for a wide range of different detailed dynamical histories. The fact that we see similar phenomenology in DNNs and in GP maps, suggests that the prediction that a local optimiser may still, on average, generate outputs with probability given by the global probability may hold more widely for highly biased input-output maps. Thus these results from DNNs support our arrival of the frequent picture in GP maps.

## S3. SUPPLEMENTARY TEXT FOR POLYOMINOES

**FIG. S5.**
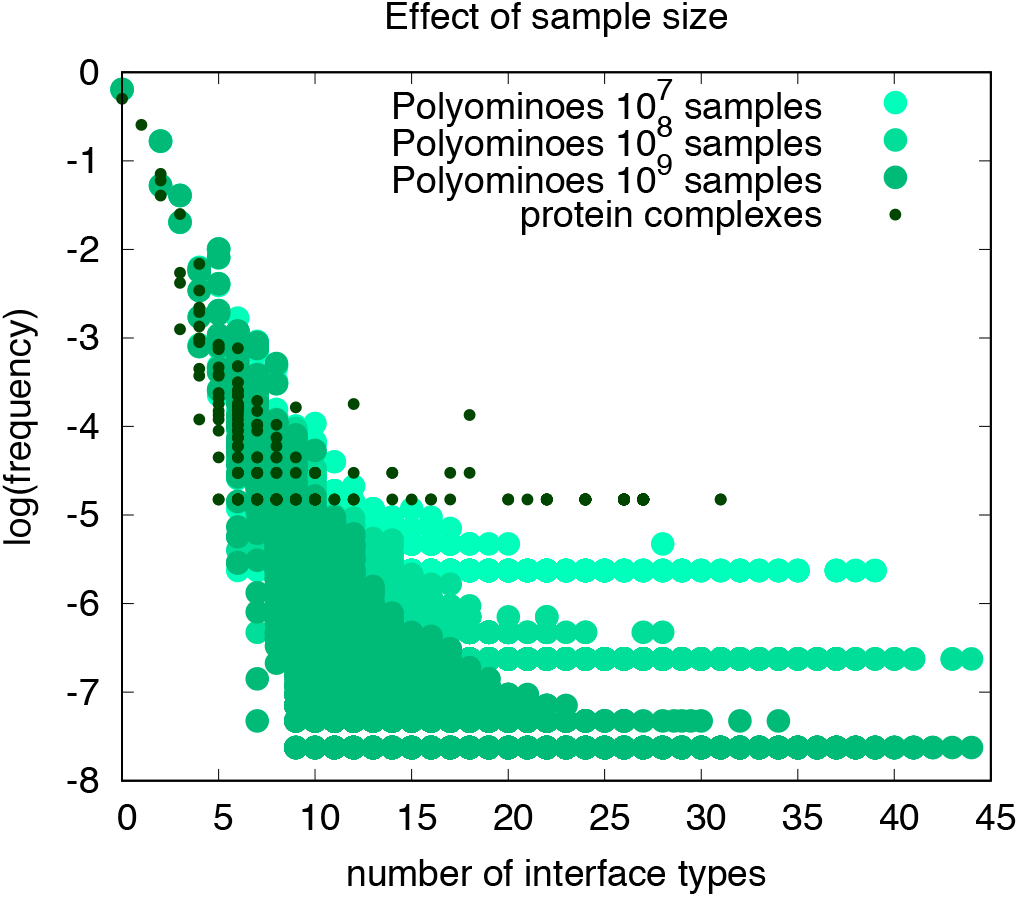
The frequency (equivalently the probability *P*(*p*)) that a polyomino *p* appears as a function of complexity 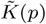= number of interfaces, for random sampling of genotypes with different numbers of samples. The plot illustrates how the long tail with high complexity is caused by finite sampling effects. In addition, the plot shows the frequency that a protein complex topology *p* is found in the PDB as a function of 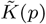, for all 34,287 topologies in the 3DComplex database (black dots). We conjecture that, just as found for polyominoes, this tail will also reduce for proteins as more topologies are published in the PDB. Nevertheless, there are likely also specific adaptive causes for certain of the low frequency/high complexity structures that may therefore have higher probability than expected from their complexity.

### A. Effects of finite sampling of genotype space on polyominoes

To test the effect of finite sampling on the *P*(*p*) versus 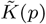 plot for polyominoes, we show in Fig. S5 data for 10^7^, 10^8^ and 10^9^ samples on *S*_4,16_, (4 tile types, 16 patch types). The main effect of a weaker sampling protocol is the emergence of a heavy tail at low frequency (*P*(*p*)) and high 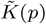. There are two reasons for this tail. The first is that we estimate 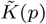 from the simplest ruleset found in the sampling that generates the polyomino, and so fewer samples means a less accurate calculation resulting in a higher complexity. However, because the simplest rulesets are typically the most likely one to be found, the error here is typically relatively small, even for structures that are rare. The second and more important reason for the long tail arises simply from poor sampling of *P*(*p*). A low *P*(*p*) structure may occur by chance just once or twice, and so be assigned a much higher frequency than if more sampling is done. This second source of error was also identified in ref. [11] (see e.g. Fig. 9 in the Supplementary Information of that paper, which shows the same tails for a circadian rhythm model where the complexities can be directly defined without sampling, so that only the second sampling argument causes the tails.). All this evidence suggests that the main source of the long tails is the poor sampling of *P*(*p*), and not measurements of 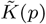.

We note that Fig. S5 also shows a large tail for the protein complex topologies. Part of the cause of this tail may also be the relatively low sampling efficiency. However, it is also likely that some rare high complexity structures are the result of specific adaptive causes. It would be very interesting to see how these trends change as more structures are deposited into the PDB.

### B. Calculating neutral space sizes in the polyomino model

In this section we analytically explore the link between the structure of a minimal genome required to produce a polyomino structure (and hence its complexity) and that structure’s neutral space space. We proceed as follows: Consider a genome *G* of length *L* that encodes a phenotype (polyomino topology) *p*. If *s* specified bonds appear in the reduced genome representation of *G*, then *l* = *L* − *s* elements of *G* are, to some extent, unconstrained. We may pick interaction labels to occupy these sites as long as these labels do not bond to the specified elements or form bonds among themselves (and thus perturb the desired structure). In this combinatoric analysis, we first calculate the number of ways of choosing a given number of labels to fill the available genome sites, so that those labels do not interfere with bonding. We then consider the ordering of these labels, and finally the orderings of the bonding information in the genome.

Consider genomes consisting of an alphabet of *n_c_* labels, *n*_0_ of which are neutral with respect to every other bond. Denote by *b* the number of bonding pairs in a reduced genome, so that *n_p_ = n_c_ n*_0_ labels remain unused in the reduced genome and bond to one other partner. We will proceed by considering the number of ways we can populate the *l* unconstrained sites in a genome with neutral labels, interacting labels that do not interfere with bonding, or a combination of both. The number of ways of picking a set of exactly *i* distinct neutral labels is

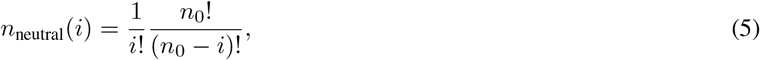

where, for later convenience, the prefactor ensures that different orderings of the set do not contribute to the total. If we choose a label that interacts with a partner to fill a neutral space, we must ensure that we do not pick the partner at any point in building our neutral string. With this constraint, the number of sets of exactly *i* bonding labels that are effectively neutral, i.e. are not able to form any bonds due to the absence of partners, is

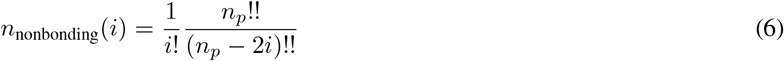

Then, the total number of ways of choosing exactly *c* distinct non-interfering labels is the sum over *i* of the ways of picking *i* neutral and *c* − *i* non-interfering but partnered labels:

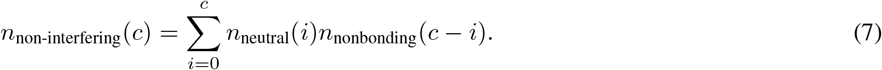

We now consider the different orderings of these labels within the available sites. The number of words of length *l* containing exactly *c* distinct characters is defined by the recursive expression

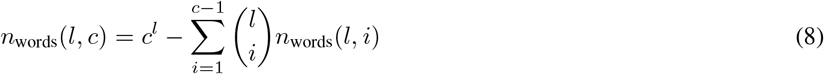

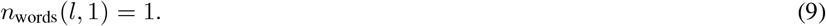

This expression equates the number of words of length *l* with exactly *c* characters to the total number of words of length *l* with at most *c* characters (*c^l^*), with all possible words with fewer than *c* characters subtracted from this total. As we ordered our sets of available colours, double counting of identical sequences is prevented. So an expression for the neutral space size consists of a combination of the above expressions for (a) ways of picking *c* distinct non-interfering colours and (b) ways of arranging these *c* colours in *l* spaces:

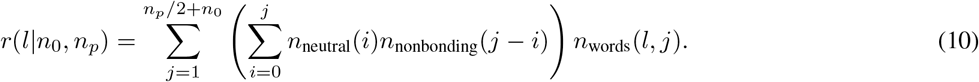

This expression gives the neutral space associated with a particular reduced genome.

As an example, consider the situation in which we have *l* = 2 spaces, *n*_0_ = 2 neutral labels, and *n_p_* = 2 unused bonding labels. If the neutral labels are `0` and `7` (as in 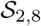), and the unused bonding labels are `5` and `6`, we can populate the empty spaces through the following:

- Use only one label to populate the two spaces (*n*_words_(2, 1)):
  - Use a neutral label: *n_neutral_*(1)*n_nonbonding_*(0)*n_words_*(2; 1) = 2 : (`00, 77`)
  - Use a bonding label: *n_neutral_*(0)*n_nonbonding_*(1)*n_words_*(2; 1) : (`55, 66`)
- Use two labels to populate the two spaces (*n*_words_(2, 2)):
  - Use only neutral labels: *n_neutral_*(2)*n_nonbonding_*(0)*n_words_*(2; 2) = 2 : (`07, 70`)
  - Use a neutral and a bonding label: *n_neutral_*(1)*n_nonbonding_*(1)*n_words_*(2; 2) = 8 : (`05, 06, 75, 76, 50, 60, 75, 76`)
  - Use only bonding labels: *n_neutral_*(0)*n_nonbonding_*(2)*n_words_*(2; 2) = 0 (as `56` would form a new bond)

So overall *r*(2|2, 2) = 14.

In general, there are also transformations to the reduced genome that leave the phenotype invariant.

1. Recalling that *b* denotes the number of bonding pairs involved in the reduced genome, the labels of bonds can be swapped for each pair (e.g. `3400 ↔ 4300`), giving

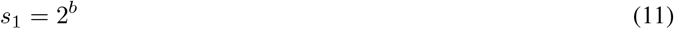

invariant genomes.
2. Any of the available bonding pairs can be used in place of any other (e.g. `3400 ↔ 5600`), giving

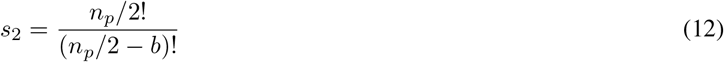

genomes.
3. Bonds can be rotated around a given subunit specification (e.g. `3400 ↔ 0340`), giving

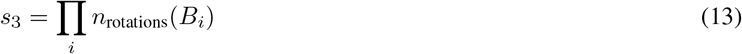

genomes, where *n*_3_(*B_i_*) is the number of invariant rotations that can be made on block *B_i_* that are not degenerate with other transformations. For example, *n*_rotations_(1111) = 1, *n*_rotations_(1112) = 4. *n*_rotations_(1122) = 2, as the arrangement `1221` cannot be reached by other transformations, but `2211` and `2112` can be reached by the pair-swapping transformation acting on the original and rotated subunit.
4. Depending on whether or not we require the seed tile for assembly to be present at a particular point in the ruleset (thus making a particular tile special), the subunits comprising the minimal ruleset may be permuted around the available space in a genome. If a ruleset consists of *n_t_* tiles, of which *t* are involved in bonding, this gives

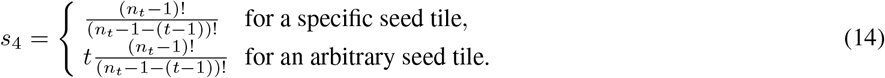
5. Finally, other symmetry-preserving transformations may be made (e.g. `1230 3400 ↔ 3120 4300`) depending on the phenotype structure: we will denote the number of transformations in this class by *s*_5_.

The overall neutral space size is then:

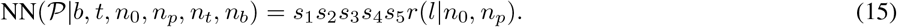

For the 2 × 1 domino in 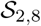 (*n*_0_ = 2; *n_c_* = 8), we have the minimal ruleset `1000 2000`, so we have *b* = 1, *n_p_* = 4, *l* = 6. Considering the above symmetric transformations that can be applied to the ruleset, we straightforwardly obtain *s*_1_ = 2, *s*_2_ = 3, *s*_3_ = 8, *s*_4_ = 1, *s*_5_ = 1. Computing *r*(6|2, 4) = 13 532, we find that the overall neutral space size is 649 536.

For the 4 × 4 16-omino in 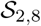, we have the minimal ruleset `1230 3400`. Properties of this ruleset immediately yield *b* = 2, *n_p_* = 2, *l* = 3. Again, consideration of the symmetry transformations above give *s*_1_ = 4, *s*_2_ = 6, *s*_3_ = 16, *s*_4_ = 1, *s*_5_ = 2. Computing *r*(3|2, 2) = 46, we find that the overall neutral space size is 35 328.

Both of these neutral space sizes are in exact agreement with those found from exhaustively sampling the 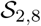 space in Ref.[26]. These examples of exact calculations of the neutral space size associated with given phenotypes illustrate the quantitative link our theory describes between algorithmic complexity and neutral space. The minimal genome for the domino structure necessitates fewer interactions than for the 16-omino (*b* = 1 and *b* = 2 respectively), leading to more degrees of freedom associated with unused bond types (*n_p_* = 4 and *n_p_* = 2 respectively). Moreover, the lower number of sites in the domino genome occupied by necessary bonding information allows more freedom associated with remaining genetic loci (*l* = 6 and *l* = 3 respectively). What this combinatorial analysis shows is that, roughly, for a linear decrease in the minimal description length, we obtain an exponential increase in the number of ways that the phenotype can be encoded within the genomes. Thus these results help illustrate how complexity and neutral space are related to one another.

Finally, we note that other combinatorial approaches have successfully predicted the neutral set sizes for RNA, as well as the difference between the rank plots of systems like RNA, which do not have redundant parts of their genomes, and systems such as the polyominoes, which do [14, 27, 28]. One potential critique of the polyomino system is that the strong phenotype bias as well as the scaling of the NSS with complexity is caused by the redundant parts of the genome which are an artefact of the fixed length encoding we use. What these more general arguments show is that the same basic phenotype bias occurs both for systems with redundant parts of the genome, and for systems which do not have this feature. See also Fig. S7 for an example of the when the genome length is changed for polyominoes.

**FIG. S6.**
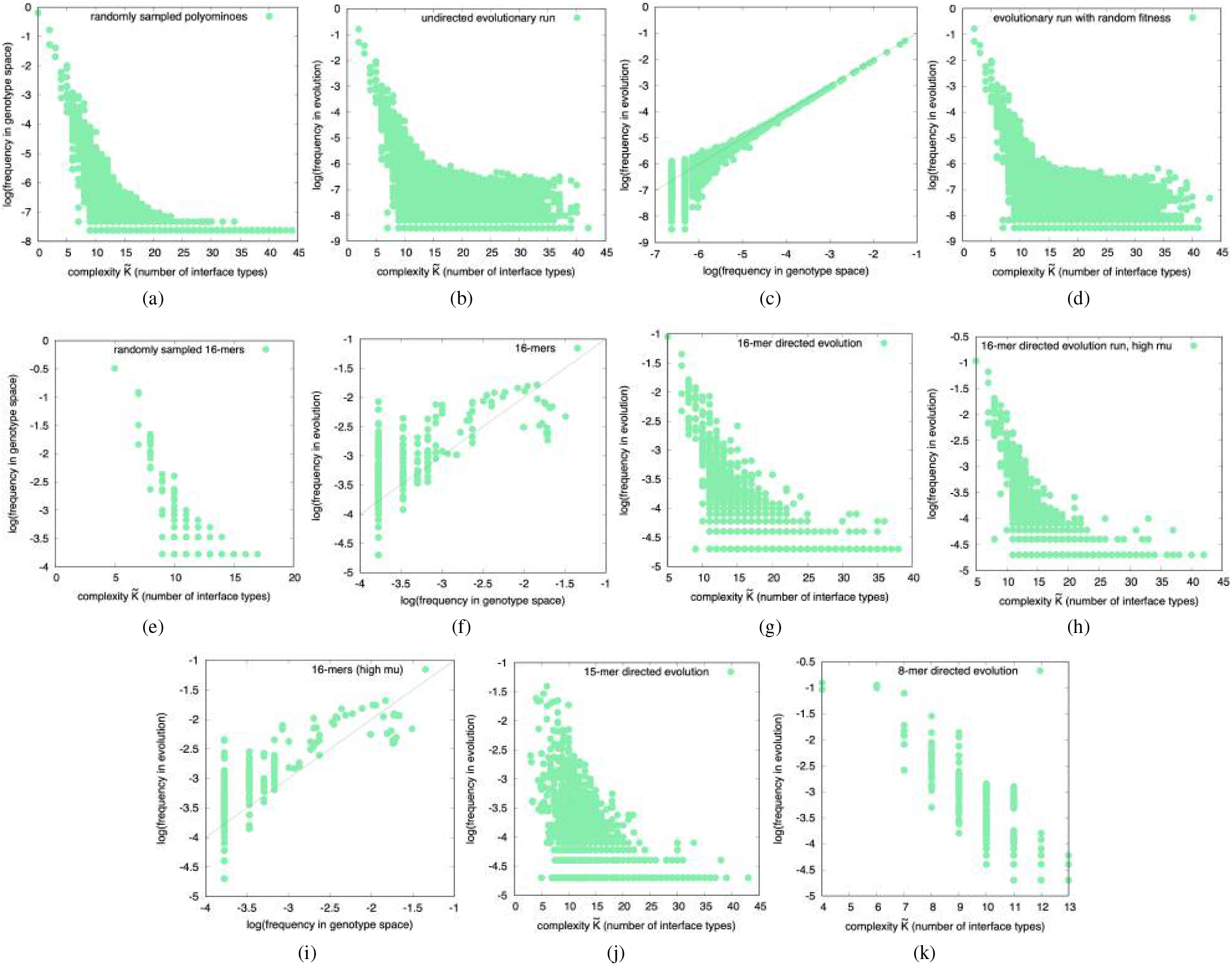
Polyomino frequency versus complexity plots for random sampling of genomes and for different evolutionary protocols. (a) The frequency (or equivalently the probability *P*(*p*) = *NSS*(*p*)*/NG*) with which (any size) polyomino structures appear upon random sampling of 10^8^ genotypes from 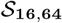 decreases with complexity 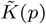 = number interface types), as expected from AIT coding theorem arguments [11]. Note that if a polyomino genome does not generate a fixed size polyomino structure deterministically, it is discarded. (b) Frequency (probability) for that a particular polyomino structure appears during undirected evolutionary simulations ( population *N* = 100 with a mutation rate *μ* = 0.1, run for 5, 000 generations) versus complexity 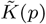 looks similar to the sampled data in (a). Here the fitness of any deterministically assembly polyomino structure was set to 1, and the fitness of non-polyomino structures (either unbounded, or structures that were not deterministic) was set to zero. (c) Frequencies from random sampling of genotypes (data from (a)) correlate well with frequencies that a phenotype arises from the undirected evolution simulations of (b). The good agreement between random sampling and the evolutionary run demonstrates that a local evolutionary population encounters new variation with a probability on average close to the global *P*(*p*) of phenotype *p* (d) Frequency (probability) that particular polyomino structure appears during versus complexity for an evolutionary run with random fitnesses protocol, where a random fitness value was applied to each polyomino shape, and reset for each evolutionary run. (e) The (normalised) frequency/probability *P*(*p*) with which 16-mer polyominoes appear upon random sampling of 10^8^ genotypes from 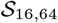 decreases with complexity 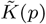, as expected from AIT coding theorem arguments [11]. Note that these 16mers are roughly one in 10^4^ of all polyominoes found in (a). (f) The probability that a 16-mer fixes in a directed evolution simulations with a fitness function with a maximum at size *s*\* = 16 for *N* = 100, mutation rate *μ* = 0.1, and run for 5000 generations, correlates well (within statistical uncertainties) with the frequency of the same16-mer structures, obtained by random sampling of genotypes. (g) Frequency (probability) for the directed evolutionary run from (f), now plotted against complexity, shows the same global bias towards low complexity shapes as for direct sampling. (h) Frequency (probability) for the directed evolutionary run for 16-mers, with *N* = 100 and a higher mutation rate *μ* = 1 is very similar to the lower mutation rate results in (f). (i) Comparing sampling of 16-mers to evolutionary simulations as in (f), but now for the high *μ* evolutionary run from (h). (j) Directed evolutionary run for 15-mers for *N* = 100 and mutation rate *μ* = 0.1 run for 5000 generations. (k) Directed evolutionary run for 8-mers for *N* = 100 and mutation rate *μ* = 0.1 run for 5000 generations. Note that for all the plots above, statistical errors are largest for the lowest frequency structures.

### C. Outcomes of evolutionary simulations with other mutation rates or fitness functions

In this section, we check that our main results for evolutionary dynamics from Fig. 1 in the main text are robust to changes in evolutionary parameters. The basics of our evolutionary simulations were described in the methods sections. Fig. S6 shows the results of a evolutionary simulations performed using a range of different evolutionary parameters and fitness functions.

In Figs. S6 (a)-(d) we study runs where any phenotype (shape) is possible, i.e. the unit fitness protocol from Sec 1B. Thus, no particular structure is most fit, and so fixes permanently. These simulations study the rate at which variation appears in an evolutionary run. Fig. S6 (a) shows the probability *P*(*p*) that a phenotype of any size appears versus complexity 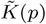 for random sampling of 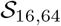 genomes (the same data can be seen in Fig. 2 of the main text). Next, in Fig. S6 (b) we show the probability-complexity relationship for evolutionary simulations where all phenotypes have the same fitness. And in Fig. S6(c), we directly compare the frequency with which phenotypes appear in the evolutionary simulation to the probability that they appear upon random sampling. As can be clearly seen, the overall probability versus complexity relationships is quite similar to that of random sampling of genotypes. There are differences at lower probabilities/higher complexities, but these may in part be due to statistical sampling issues. Note that the *y*-axis is on a log-scale, so that these phenotypes are found rarely, and that the majority of the probability weight is in the regime where the data from (a) and (b) are very similar. As argued above, and also in refs. [6, 8], the mean probability that a phenotype *p* appears in a population during an evolutionary run, which is fundamentally measuring a *loca*l quantity, is very well approximated by the *global* frequency *P*(*p*). When this phenomenology is at play, then simple analytic forms such as those shown in Section S2 work well. Finally, in Fig. S6 (d) we show the outcome of simulations where where for each run, each bound, deterministic polyomino structure is assigned a fitness value uniformly distributed on [0, 1], i.e. the *random fitness* protocol from Methods. Again, as expected within the arrival of the frequent picture, the probability-complexity relationships are remarkably similar to that obtained by random sampling of genotypes, and are also very close to the outcomes of the unit fitness protocol.

In Figs. S6(e)-(i), we show results for simulations and sampling with a focus on polyominoes of size *s* = 16, using the *size fitness* protocol from methods, where the fitness function for a polyomino of size *s* is 1/( |*s* − 16| + 1), so that polyominoes of size 16 have unit fitness and other sizes have fitness decreasing with distance from 16. First, in Fig. S6(e) we show the probability *P*(*p*) that a 16-mer appears upon random sampling of 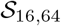 genomes vs. complexity 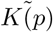. As expected from the coding theorem bound Eq. (17) (Eq.(1) in the main text), we once again see behaviour consistent with an upper bound that decreases exponentially for linear increases in complexity. Next, in Fig. S6(g) we compare the frequency with which 16-mers fix in an evolutionary simulation, to the probability that they appear upon random sampling (this data can also be seen in the inset of Fig. 2 of the main text). Although there is less data, so that the fluctuations are larger, we observe the same linear relation as seen in Fig. S6 (c). Fig. S6(g) directly shows the evolutionary data for the fixation probability of 16-mers, which closely resembles the sampled data from Fig. S6(e). In Fig. S6(h) we show data for simulations with a much higher mutation rate *μ*, and finally, in Fig. S6(i) we directly compare the frequency with which 16-mers appear in an evolutionary simulation, to the probability that they appear upon random sampling. Again, as expected from predictions from Eqs.(1) and (2), the data is very similar to that found with the lower mutation rate, in fact the frequency versus complexity is slightly closer to the sampling results. This closer agreement may be due to the high *μ*: if evolutionary search is more randomised, the global effect of *NSS* on evolutionary observation will become ever stronger. Conversely, at low *μ*, evolution is more locally constrained, and effects from the local structure of search space can act to drive the relationship away from simple global scaling with *P*(*p*). But more work is needed to confirm this conjecture.

The final two plots Figs. S6(j) and (k), show the frequency versus complexity relationship for evolutionary runs for fitness functions *f* = 1/( |*s* − 15| + 1), which favours 15-mers, and *f* = 1/(|*s* − 8| + 1), which favours 8-mers. Both experiments show qualitatively similar behaviour to the runs for 16-mers.

Overall these results corroborate our claims in the main text, which is that he *arrival of the frequent* means that a highly biased GP map strongly constrains what variation can arise in practice, and that this results in evolutionary outcomes that strongly favour low complexity phenotypes over high complexity phenotypes. While this is the broad overall picture, a closer look at Fig. S6 suggests many more subtle effects which would be interesting to explore in the future.

### D. Symmetry and modularity in evolved polyominoes

#### 1. Enumeration of polyomino symmetries

The distribution of symmetries of polyominoes have been characterised up to size *n* = 27. These data are available from the On-Line Encyclopedia of Integer Sequences, http://oeis.org/ and are curated and tabulated in Table I.

**TABLE I.** Symmetries of polyominoes of different sizes. (a) Symmetry groups for polyominoes of size *n*. (b) Symmetry groups for all polyominoes of size up to and including *n*. Data taken from the On-Line Encyclopaedia of Integer Sequences, http://oeis.org/ under the following references: (all polyominoes) A000105; (*D*_4_) A142886; (*C*_4_) A144553; (*D*_2_) A056877 + A056878 (diagonal and axial symmetry); (*C*_2_) A006747; (*D*_1_) A006746 + A006748 (diagonal and axial symmetry); (*C*_1_) A006479.

| n | (a) $D_4$ | $C_4$ | $D_2$ | $C_2$ | $D_1$ | $C_1$ | (b) $D_4$ | $C_4$ | $D_2$ | $C_2$ | $D_1$ | $C_1$ |
| --- | --- | --- | --- | --- | --- | --- | --- | --- | --- | --- | --- | --- |
| 1 | 1 | 0 | 0 | 0 | 0 | 0 | 1 | 0 | 0 | 0 | 0 | 0 |
| 2 | 0 | 0 | 1 | 0 | 0 | 0 | 1 | 0 | 1 | 0 | 0 | 0 |
| 3 | 0 | 0 | 1 | 0 | 1 | 0 | 1 | 0 | 2 | 0 | 1 | 0 |
| 4 | 1 | 0 | 1 | 1 | 1 | 1 | 2 | 0 | 3 | 1 | 2 | 1 |
| 5 | 1 | 0 | 1 | 1 | 4 | 5 | 3 | 0 | 4 | 2 | 6 | 6 |
| 6 | 0 | 0 | 2 | 5 | 8 | 20 | 3 | 0 | 6 | 7 | 14 | 26 |
| 7 | 0 | 0 | 4 | 4 | 16 | 84 | 3 | 0 | 10 | 11 | 30 | 110 |
| 8 | 1 | 1 | 5 | 18 | 28 | 316 | 4 | 1 | 15 | 29 | 58 | 426 |
| 9 | 2 | 0 | 4 | 19 | 64 | 1196 | 6 | 1 | 19 | 48 | 122 | 1622 |
| 10 | 0 | 0 | 9 | 73 | 112 | 4461 | 6 | 1 | 28 | 121 | 234 | 6083 |
| 11 | 0 | 0 | 12 | 73 | 238 | 16750 | 6 | 1 | 40 | 194 | 472 | 22833 |
| 12 | 3 | 3 | 18 | 278 | 420 | 62878 | 9 | 4 | 58 | 472 | 892 | 85711 |
| 13 | 2 | 2 | 20 | 283 | 890 | 237394 | 11 | 6 | 78 | 755 | 1782 | 323105 |
| 14 | 0 | 0 | 35 | 1076 | 1595 | 899265 | 11 | 6 | 113 | 1831 | 3377 | 1222370 |
| 15 | 0 | 0 | 41 | 1090 | 3334 | 3422111 | 11 | 6 | 154 | 2921 | 6711 | 4644481 |
| 16 | 5 | 12 | 74 | 4125 | 6013 | 13069026 | 16 | 18 | 228 | 7046 | 12724 | 17713507 |
| 17 | 4 | 7 | 73 | 4183 | 12547 | 50091095 | 20 | 25 | 301 | 11229 | 25271 | 67804602 |
| 18 | 0 | 0 | 137 | 15939 | 22824 | 192583152 | 20 | 25 | 438 | 27168 | 48095 | 260387754 |
| 19 | 0 | 0 | 148 | 16105 | 47468 | 742560511 | 20 | 25 | 586 | 43273 | 95563 | 1002948265 |
| 20 | 12 | 44 | 292 | 61628 | 86832 | 2870523142 | 32 | 69 | 878 | 104901 | 182395 | 3873471407 |
| 21 | 7 | 25 | 273 | 62170 | 180531 | 11122817672 | 39 | 94 | 1151 | 167071 | 362926 | 14996289079 |
| 22 | 0 | 0 | 539 | 239388 | 332010 | 43191285751 | 39 | 94 | 1690 | 406459 | 694936 | 58187574830 |
| 23 | 0 | 0 | 540 | 240907 | 689858 | 1.68046E+11 | 39 | 94 | 2230 | 647366 | 1384794 | 2.26234E+11 |
| 24 | 20 | 165 | 1161 | 932230 | 1273985 | 6.54997E+11 | 59 | 259 | 3391 | 1579596 | 2658779 | 8.81231E+11 |
| 25 | 11 | 90 | 1026 | 936447 | 2647385 | 2.55722E+12 | 70 | 349 | 4417 | 2516043 | 5306164 | 3.43845E+12 |
| 26 | 0 | 0 | 2128 | 3641945 | 4907236 | 9.99908E+12 | 70 | 349 | 6545 | 6157988 | 10213400 | 1.34375E+13 |
| 27 | 0 | 0 | 2006 | 3651618 | 10197786 | 3.9153E+13 | 70 | 349 | 8551 | 9809606 | 20411186 | 5.25905E+13 |

**FIG. S7.**
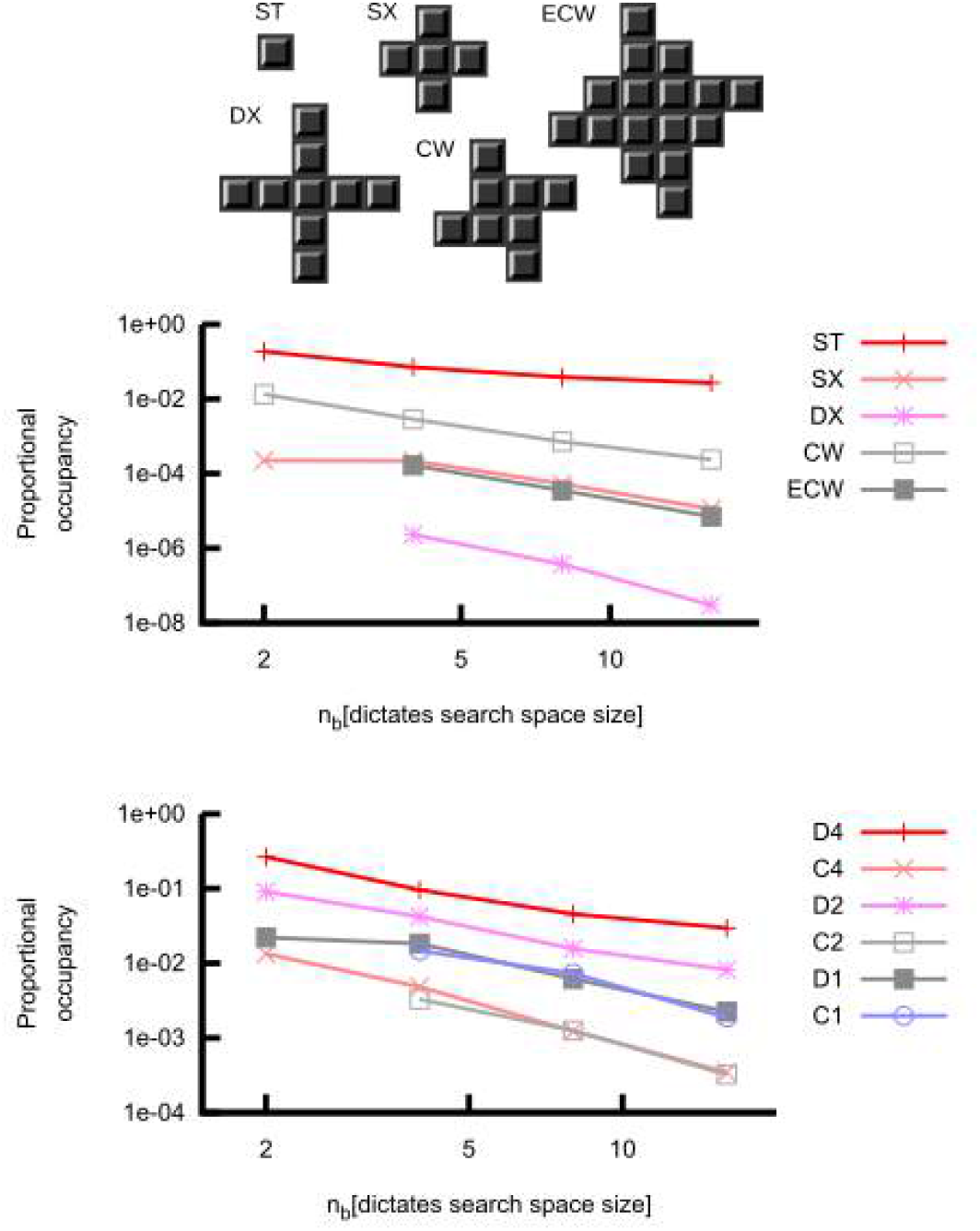
Proportion of genome space that encodes for a particular tiles or symmetry group, versus the number of tiles n_b_ in spaces of the form 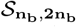 **ranging from** 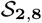 **to** 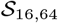. Top figure: some key tile shapes. Middle figure, data denotes the frequency of individual tiles shown in the top figure. Bottom, the data are for all tiles of symmetries *D*_4_, *C*_4_, *D*_2_, *C*_2_, *D*_1_ and *C*_1_. Increasing the size of the space allows for more diversity in shapes, and in symmetries, as can also be seen in table 1. As *n_b_* increases, the fraction of genomes leading to non-deterministic or unbounded structures also increases; this explains why the overall occupancies of the different polyomino structures decreases for larger spaces.

**FIG. S8.**
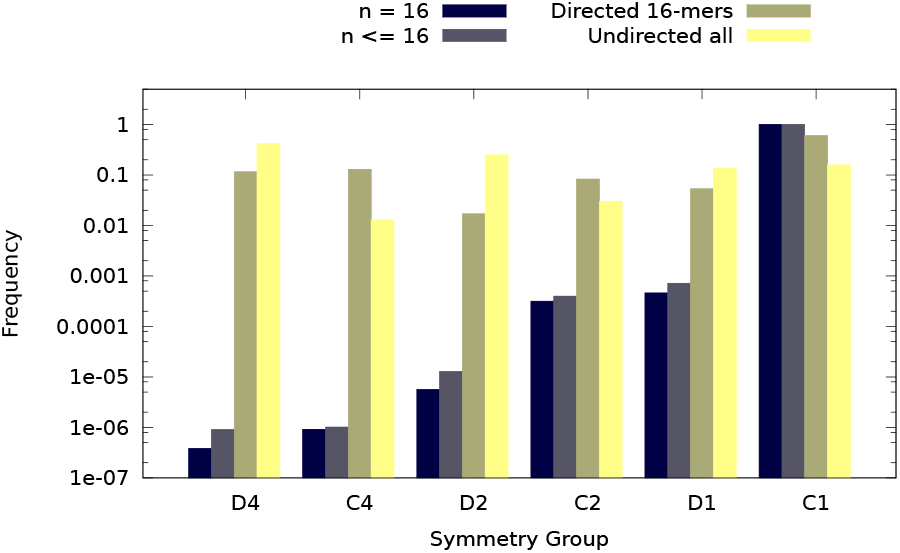
Comparison of symmetry groups for undirected evolutionary run and a run directed towards 16-mers. The two evolutionary simulations are for *N* = 100, *μ* = 0.1 and run for 50,000 generations on 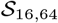. In one run, all finite self-assembling polyominoes have a fitness 1, and in the other directed evolutionary simulation a fitness maximum at size *s*\* = 16 is used. The frequencies for each symmetry group, found by the evolutionary simulations are compared to the overall frequencies from table I for *n* = 16 (black), and *n* ≤ 16 (brown). The evolutionary simulations are very strongly biased towards high symmetry structures.

**FIG. S9.**
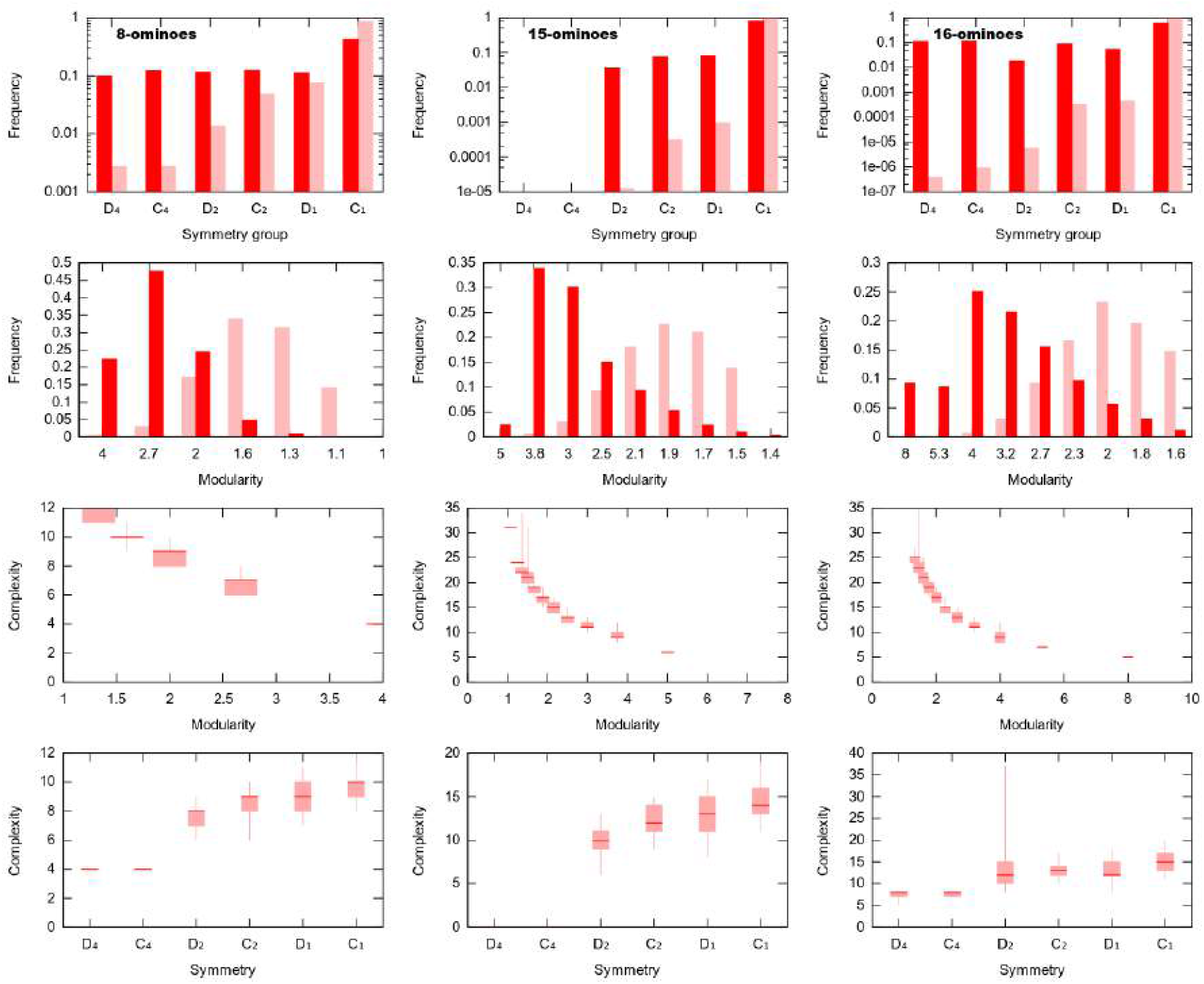
Symmetry, modularity and complexity in evolved polyomino structures. Each column presents results of polyomino structures resulting from evolutionary simulation directed towards *s*\* = 8, 15, 16. (Row 1) Symmetry classes observed in evolved polyominoes (dark) compared to across all polyominoes of that size (light), showing a dramatic favouring of symmetric structures. (Row 2) Modularity index *S/b* observed in evolved structures, showing a favouring of higher-modularity structures up to a maximum index of *s*\*/2. (Row 3) Distribution of 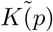 complexity values (number of interface types) among polyominoes of a given modularity index shows a clear decrease of complexity with increasing modularity. Candlesticks show 0.05, 0.25, 0.5, 0.75, 0.95 quantiles. (Row 4) Distribution of 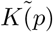 complexity values among polyominoes of a given symmetry class shows a clear decrease of complexity with increasing symmetry. Candlesticks show 0.05, 0.25, 0.5, 0.75, 0.95 quantiles.

#### 2. Scaling of symmetry group proportions with search space size

How do our results hold across varying search space sizes? This is a consideration when interpreting our results in a biological context, as the genetic search spaces associated with organisms in nature are not of fixed size. For example, gene duplications can vary the length of genomes. We explored the proportion of several differently sized search spaces occupied by genomes encoding specific polyomino structures, and the proportion occupied by genomes encoding structures that fall within various symmetry groups. Fig. S7 shows these proportions for search spaces ranging 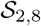 to 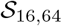, covering many orders of magnitude in absolute search space size. As search space grows, we observe a general decrease in the proportion of search space encoding any given structure, which is unsurprising, as larger search spaces support more diverse structures. To first order, the proportion of high symmetry structures encoded by the genomes does not vary so much over this large variation in the size of the spaces.

#### 3. Symmetry and modularity of polyominoes from evolutionary simulation

In Fig. S8 we display specific structures and symmetry classes observed in evolutionary simulation of polyomino structures, both undirected and directed towards *s*\* = 16. As can be easily seen, in both cases there is a very strong bias towards low complexity/high symmetry structures. The undirected runs have slightly more higher symmetry structures because a higher proportion of the smaller structures have higher symmetries (compare Table I (a) and (b)).

Next, in Figs. S9 (a)-(c) we compare the symmetries under evolutionary runs from above, to those from Table I(a) for 8-mers, 15-mers, and 16-mers. In each case we observe a strong bias towards symmetric structures.

In the following row, Figs. S9 (d)-(f) we compare a modularity index which is defined as the size *s* of the polyomino divided by the number of tile types *n_b_* used in a minimal genome. This index measures how often units are repeated. For example, a modularity of 1 means every tile is different, whereas a modularity of *s/*2 means that just two tiles are used, on average *s/*2 times. Thus a higher modularity index means that the system is built out of a smaller number of modular units; one could say it is more modular (although biological modularity is a much more complex feature than just something that is repeated [29, 30]). As can be clearly seen in Figs. S9 (d)-(f), the evolutionary simulations are biased towards structures with a higher modularity index. We did not work out the modularity index of all structures, as that would require calculating their minimal rule sets which is prohibitively expensive. Instead, we took all rule-sets found in the simulation, and counted each structure once to obtain a distribution of modularity indices. If we had, instead, taken all structures for each size, then the bias towards high modularity would be (much) more pronounced.

In the next row, Figs. S9 (g)-(i), we compare our modularity index to complexity. As expected, the complexity increases with increasing modularity index. Note that the complexities are calculated for those structures that appear in the evolutionary simulation, which has a bias towards low complexity. In the final row, Figs. S9 (j)-(l), we compare symmetries to our complexity measure 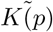. As expected, the complexity increases with decreasing symmetry. Note that for these final two rows, the complexities are calculated for those structures that appear in the evolutionary simulation, which has a bias towards low complexity. If we were to compare the complexity and symmetry by uniformly sampling over all structures, then there would be more variation in the complexity.

**FIG. S10.**
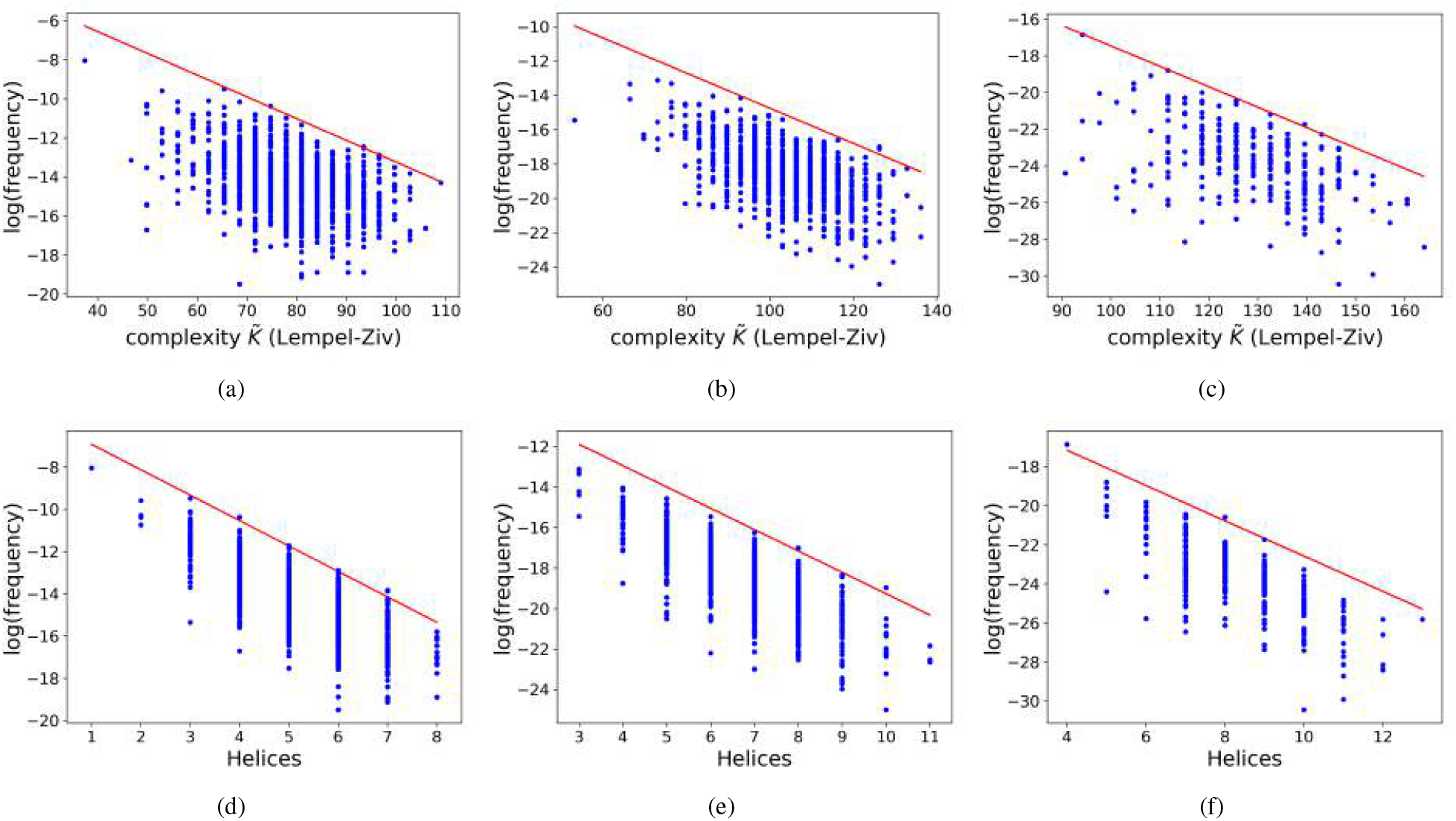
Frequency (calculated using the NNSE neutral set size estimator [31]) versus complexity for natural RNA SS from the fRNAdb, for lengths *L* = 75, *L* = 100, and *L* = 126. There are 1419 sequences for *L* = 75, 932 for *L* = 100 and 318 for *L* = 126. The red line is an upper bound, Eq. 17 (Eq. (1) of the main text) consistent with the data. Parameters *a* and *b* are fit. Complexity is measured in two ways: by *CLZ* from Eq. (18) for (a) *L* = 75, *a* = 0.37 and *b* = 7; (b) *L* = 100, *a* = 0.34 and *b* = 15; and (c) *L* = 126, *a* = 0.37 and *b* = 21; and by the number of helices for (d) *L* = 75, *a* = 4 and *b* = 19; (e) *L* = 100, *a* = 3.5 and *b* = 29; (f) *L* = 126, *a* = 3 and *b* = 45. Both measures show the same overall scaling. The top probability structure for each value of the *C_LZ_* complexity for *L* = 100 natural data in (b) is also shown in table III.

**FIG. S11.**
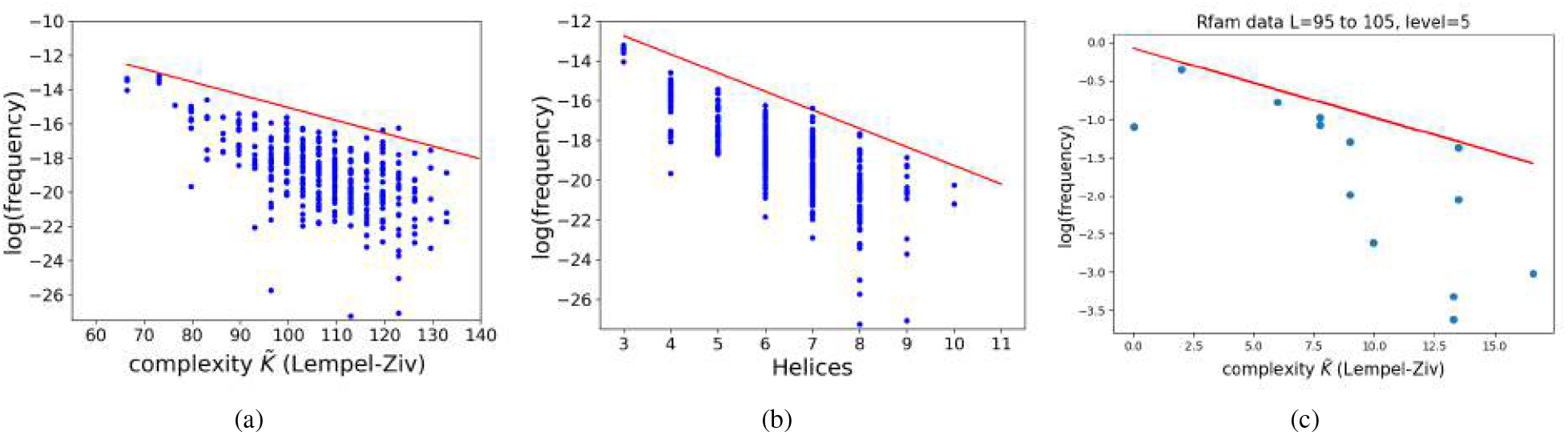
Frequency versus complexity for natural RNA structures from the Rfam database [32, 33]. (a) *L* = 100 using the NNSE neutral set size estimator [31] to estimate frequencies versus complexity (measured by *C_LZ_* from Eq. (18)) for natural RNA; The red line is and upper bound, Eq. 17 (Eq. (1) of the main text) consistent with the data. Parameters *a* = 0.25 and *b* = 25 are fit. (b) the same data as (a) but with complexity measured as number of helices (*a* = 3.1 and *b* = 33); (c) Level 5 coarse-grained structures for *L* ≈ 100 (lengths *L* = 95 to 105 were binned together improve the reliability of database frequency estimates). There are 4124 sequences. As opposed to the other graphs in this section, the frequencies are directly taken from the Rfam database. Complexity is measured by *C_LZ_*, but the structures come from consensus sequence alignment methods, not from folding calculations. The bound of Eq. (18) is for *a* = 0.3 and *b* = 0.25. The data is similar to that extracted from the fRNAdb in Fig. 3(b) of the main text. For all three Rfam data graphs, the same overall scaling of complexity with probability is observed as for the *L* = 100 data from the fRNAdb.

## S4. SUPPLEMENTARY TEXT FOR RNA GP MAP

### A. Probability-complexity for natural RNA from fRNAdb and Rfam with frequencies calculated with the NNSE

In Fig. S10 we plot frequency versus complexity 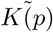 for natural RNA. The structures are computationally predicted from natural sequences from the fRNA database, and the probabilities are estimated using the NNSE from [31] for each sequence in the database. This method is the one that was used in ref. [16], and it is indirect. By contrast the frequencies/ probabilities in the main text are directly sampled. Here we show the probability versus complexity measured firstly in (a)-(c) with the standard Lempel-Ziv measure from ref. [16], shown in Eq. (18), and secondly, for (d)-(f) for a simpler measure, namely the number of helices or stacks, which has been proposed before as a complexity measure [34], and which also shows similar scaling to that expected from the upper bound of Eq. (17) (or equivalently, Eq. (1) in the main text). One can observe that the scaling for these longer lengths looks cleaner than in the main text for *L* = 30, where finite size effects play a more important role.

As a quick check that database biases and structure prediction methods are not the main causal factor in our observations, in Fig. S11 we make probability-complexity plots for RNA with *L* ≈ 100 data from the Rfam database [32, 33]. The figure shows analogous plots to Fig. S10, with frequencies estimated via the NNSE [31], and complexity via *C_LZ_* and via the number of helices.

Fig. S11(c) a frequency-complexity plot where both the structures and the frequencies are derived via the same methods as in Fig. 3 of the main text, which is different from the NNSE method used in Fig. S10: The frequencies are simply direct database frequencies of abstract shapes (level 5), and the secondary structures are obtained via consensus sequence alignment methods, which is fundamentally different from the energy minimization method used elsewhere in this work. The structures were taken from all available seed sequences of ncRNA families from the Rfam database (but some of the structures were ignored due to containing impossible/unusual motifs such as loops of zero length). We used lengths 95 to 105 and binned these together due to a paucity of data. Just as in the main text Fig. 3, abstract level 5 was used. It is evident that the same general relation of probability to complexity is observed as for the fRNAdb. This similarity is further support for the claim that the probability-complexity relationships are not due to database artefacts.

In Table II we show how our Lempel-Ziv complexity measure *C_LZ_* (*x*) from Eq(1) in the main text, or Eq. (18) in these SM, correlates with the probability *P*(*p*) that a particular RNA SS phenotype *p* obtains upon random sampling of sequences. The table also shows the dot-bracket notation of the SS, which helps illustrate what *C_LZ_* (*x*) is measuring. In Table III we show a similar plot, but now for natural *L* = 100 RNA from the fRNAdb, which are also shown in Fig. S10(b). Note that, as expected, the range of complexities in these natural RNAs is smaller than the fully sampled ones for *L* = 55 in Table II.

**TABLE II.**
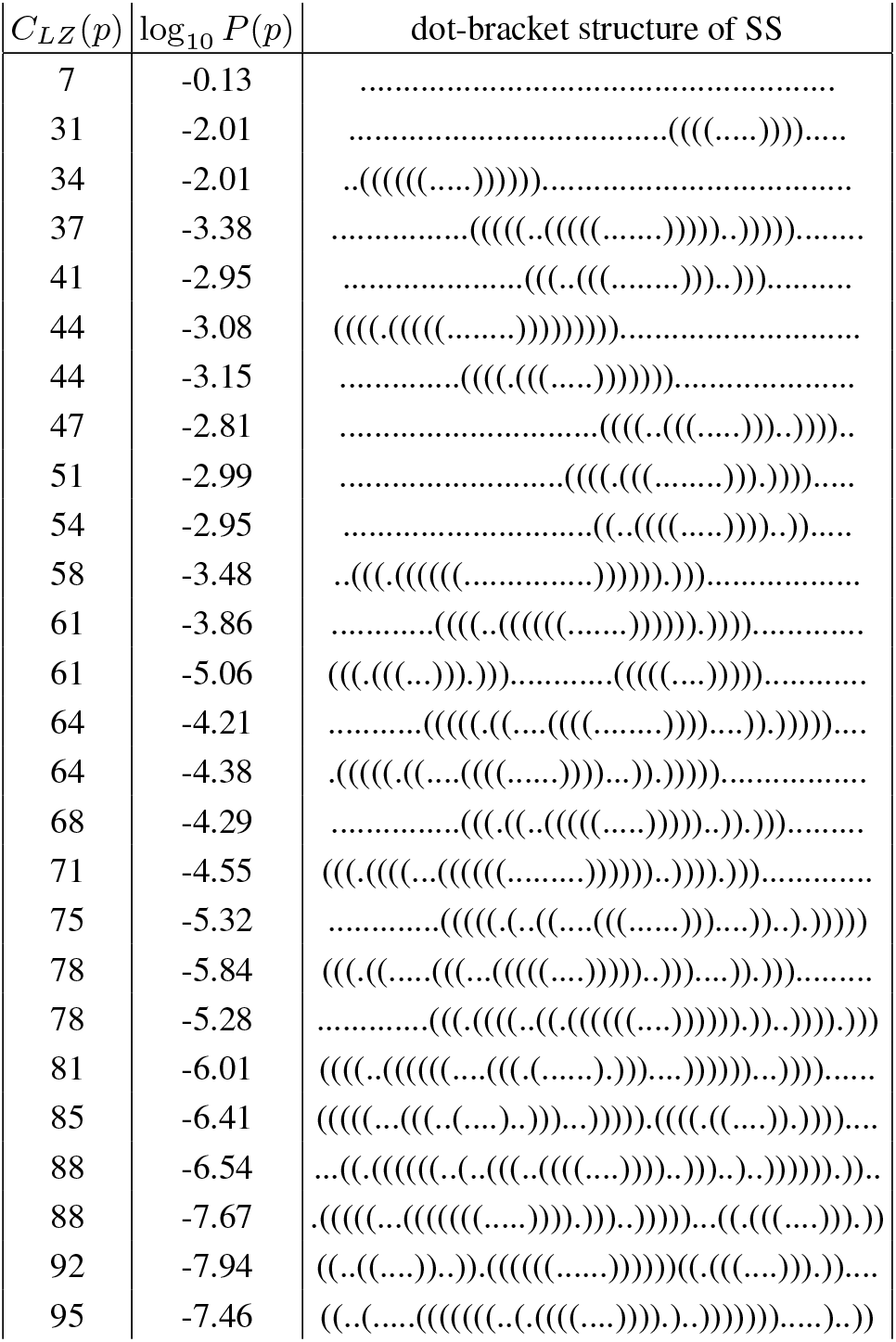
An example of how complexity 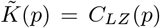, probability *P*(*p*) and the dot-bracket description of secondary structures for *L* = 55 RNA relate to one another. Structures were generated with the Vienna package [35] upon random sampling of sequences, and the probabilities were calculated with the NSSE [31]. For each complexity, the highest probability value SS was chosen. A clear decrease of probability with increasing complexity can be observed, which is consistent with Eq. (17).

**TABLE III.**
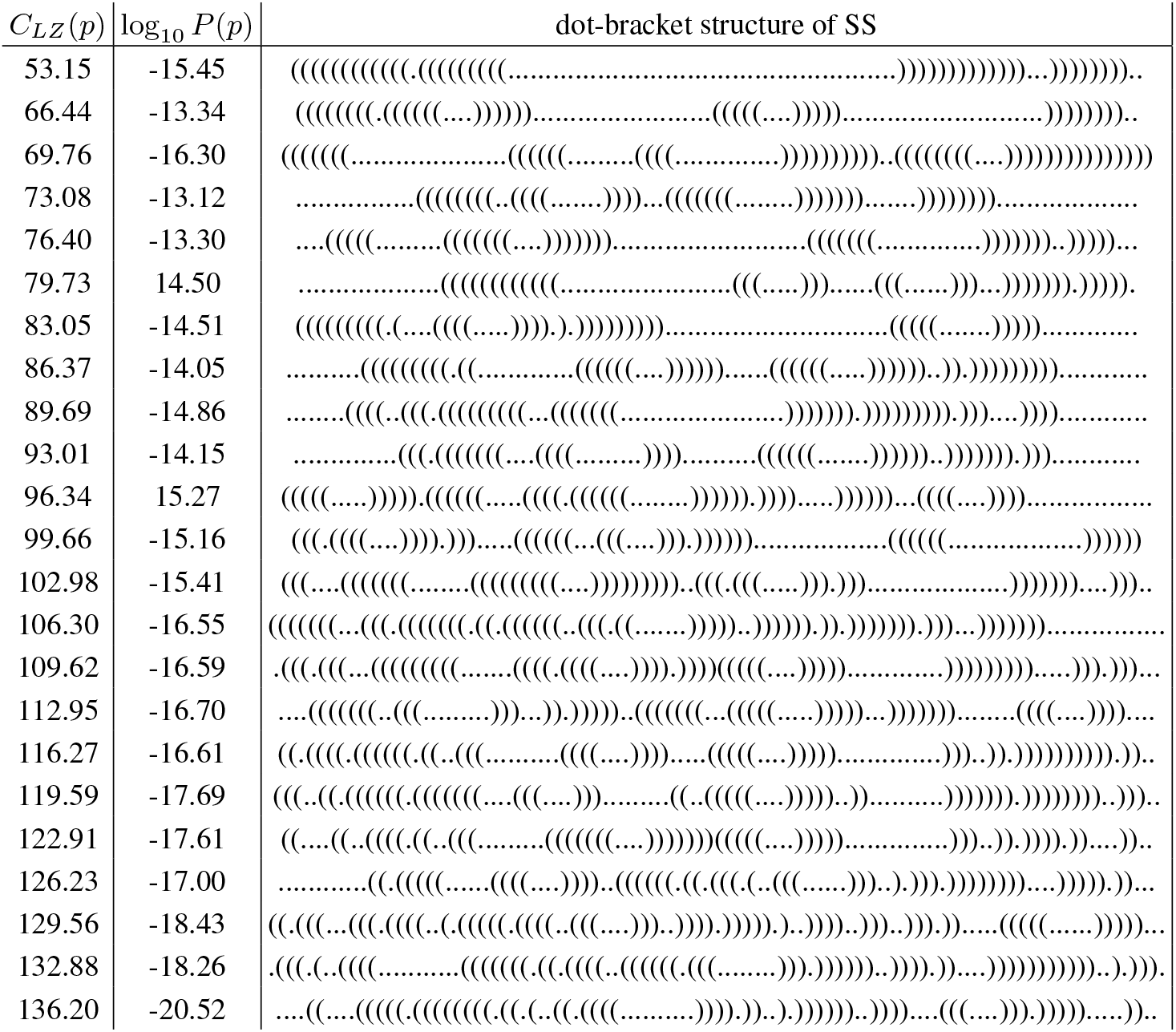
An example of how complexity 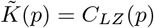, database frequency *P*(*p*) and the dot-bracket description of secondary structures for naturally occurring *L* = 100 RNA relate to one another. The structures are from fRNAdb, and for each one, the probabilities were calculated with the NSSE [31]. For each complexity, the highest probability value SS was chosen. The full set of probabilities and complexities can be seen in Fig. S10(b). A clear decrease of probability with increasing complexity can be observed, as expected from Eq. (17) and as can also be seen in Fig. S10(b)

### B. RNA mutational robustness

The mutational robustness of a given structure *p* is defined as the fraction of random single point mutations of a random sequence underlying *p*, which map to *p*. The mutational robustness of RNA has been studied previously, and it is known that the robustness of a structure scales roughly linearly with the log probability of the structure [8, 14, 28]. Here we provide some further evidence for this scaling. Fig. S12(a) shows a roughly linear relation between mutational robustness and log frequency for *L* = 18. To make the plot, 10^6^ samples were chosen, and all 3*L* single point mutations were enumerated, and each of these mutated sequences were folded to their associated structures. The linear relation is more clear for higher probability phenotypes. *L* = 18 was chosen as a small enough system that the linear relation becomes apparent, even on partial sampling. Next, Figs S12(b) and (c) show the same approximately linear relation, but now for *L* = 30 with 10^6^ randomly sampled sequences and natural RNA data, respectively. There is more spread, in part because of the much larger number of sequences and structures, but to first order this scaling still holds.

We showed earlier [16] also that for *L* = 55 ncRNA structures, the distribution of random and natural robustness values were virtually identical, while both being very different to the distribution obtained via directly sampling phenotypes (P-sampling). In Fig. S12(d), the directly calculated distribution of robustness values for *L* = 30 natural data and randomly sampled structures (G-sampled) are shown. Additionally, an estimate of the robustness distribution over all phenotypes (P-sampling) is given in black, which is obtained using the same methods described in ref. [16]. Those calculations make use of accurate estimates for the log neutral set size distribution over all phenotypes, combined with a rough fit to the robustness-log probability plots for *L* = 30. Firstly, the natural and random distributions clearly differ significantly from the estimated P-sampled distribution over all structures. Secondly, the random and natural distributions are remarkably similar. Interestingly, in contrast to the case found in [16] for *L* = 55, where we found no statistical difference between natural data and randomly sampled data, for *L* = 30 we observe that the natural data has slightly higher mean robustness values: 0.36 for the natural data vs. 0.32 for the random samples. While this difference in means is statistically significant (two sample t-test *p*-value*<* 10^−100^), the absolute difference is very small. Nevertheless, it suggests that there may have been a small amount of natural selection for higher robustness in these shorter strands. The P-sampled mean robustness is 0.18, considerably less than either the random or natural mean robustness values.

One possible alternative explanation for the overall correlation we find between frequency and neutral set size is that there exists a strong selection for robustness. While it is true that a selection for robustness would lead to deviations from P-sampling, itwould be extremely surprising if this led to the overall distribution of robustness being so close to the randomly G-sampled one, as seen for *L* = 55 [16] and *L* = 30 structures here. Thus this potential alternative hypothesis is highly unlikely to hold.

**FIG. S12.**
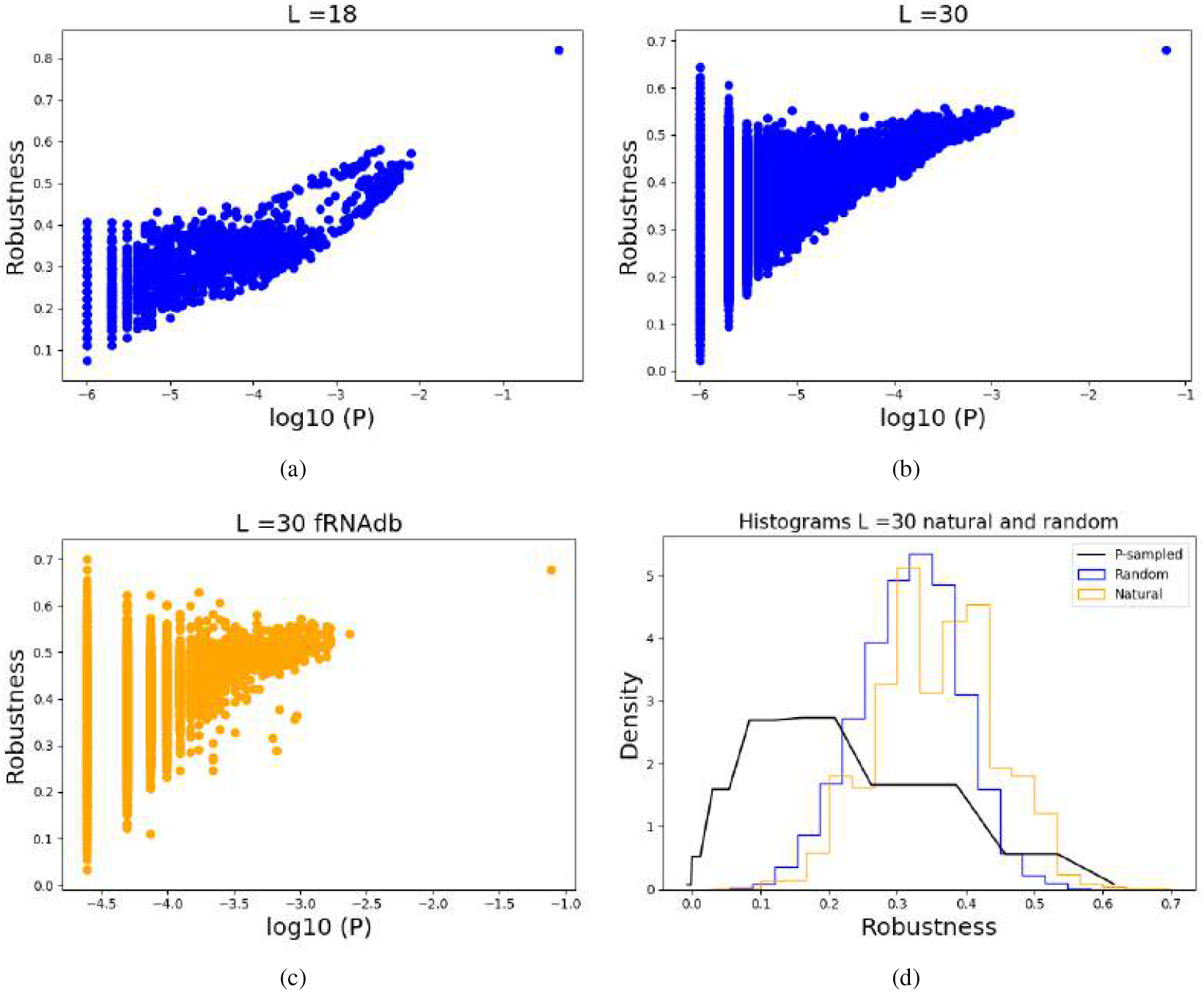
RNA mutational robustness plots: (a) A roughly linear relation between robustness and log probability for randomly sampled RNA, *L* = 18; (b) A roughly linear relation between robustness and log probability for randomly sampled RNA, *L* = 30 and (c) natural fRNAdb *L* = 30 data; (d) Natural and random RNA *L* = 30 have very similar robustness distributions, and markedly different from the P-sampled (direct phenotype sampled) distribution. A very similar close correlation between randomly sampled structures and natural structures was shown for *L* = 55 [16].

## S5. SUPPLEMENTARY TEXT FOR ALGORITHMIC INFORMATION THEORY AND COMPLEXITY MEASURES

### A. Algorithmic information theory, Kolmogorov complexity and the AIT coding theorem

The word *complexity* is used in many different ways across science (see e.g. ref. [36] for an overview), and this wide variation in meaning often causes confusion. In biological contexts measures of complexity related to Shannon information theory have been widely applied [34, 37–39]. An alternative approach is to use algorithmic information theory (AIT), which is a different concept, although there are deep links between Shannon information and AIT [40]. Here we focus on measures related to Kolmogorov complexity, a key concept from AIT, which formally captures the intuition that strings with short descriptions have lower complexity than strings that only allow long descriptions. Consider, for example the two 100 digit long binary strings below:

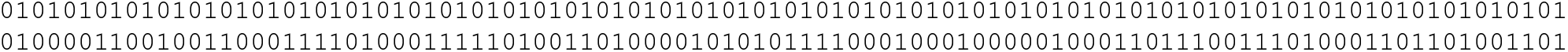

The first string intuitively has low Kolmogorov complexity because it can be described by a small program such as “print “01” 50 times”, while the second string has, as far as we know, no shorter program than to simply print the full string. More formally, Kolmogorov complexity *K*(*x*) is defined as the length of the shortest program that generates a string *x* on a suitably chosen (optimal) universal Turing machine (UTM). Since a universal Turing machine can mimic any other Turing machine, there is an *invariance theorem* [40, 41] which states that if *K_U_* (*x*) and *K_V_* (*x*) are the Kolmogorov complexities defined w.r.t UTMs *U* and *V* respectively, then |*K_U_* (*x*) − *K_V_* (*x*)| ≤ *M_U,V_*, where *M_U,V_* is a constant independent of *p*. The actual UTM used is then often informally dropped, and so we speak simply of *K*(*x*). Nevertheless, it is important to remember that Kolmogorov complexity is always defined w.r.t. a particular UTM (See ref. [40] for more in-depth discussion.).

An important link between Kolmogorov complexity and probability was derived by Levin [42], a result which is related to earlier work on algorithmic probability by Solomonoff [43] who in fact provided the first formulation of what is now called Kolmogorov or Kolmogorov-Chaitin complexity. Very briefly ( for more technical details, see e.g. ref. [40]) the AIT coding theorem states that the probability *P* (*x*) that a randomly selected (binary) input program fed into a (optimal prefix) universal Turing machine (UTM) generates output *p* is bounded by

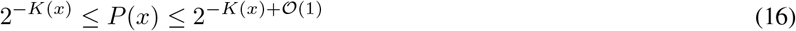

where *K*(*x*) is the Kolmogorov complexity of output *x*. The (unknown) 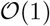 term is independent of *x*, and hence in an asymptotic regime of large complexities, can be ignored. Informally, what Eq. (16) tells us is that upon randomly choosing programs, a UTM is exponentially more likely to produce outputs with low *K*(*x*) than outputs with high *K*(*x*).

Despite its profound implications, the coding theorem has not been widely applied in science and engineering. One of the reasons is that many practical systems are not UTMs, and the coding theorem (like much of AIT) depends on the computational power of UTMs. The second reason is that the Kolmogorov complexity *K*(*x*) is formally uncomputable due to the famous halting problem of UTMs, first pointed out by Turing [44]. In a recent paper [11] that was inspired by the AIT coding theorem, a practically useable upper bound for the *P* (*x*) that output *x* obtains upon uniform sampling of inputs was derived for input-output maps *f* : *I* → *O* that are *simple*, that is *K*(*f*) is independent, or grows very slowly with the size of the output space. It takes the form

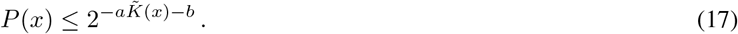

where 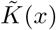 is a suitable approximation to the Kolmogorov complexity of output *x*, and *a* and *b* are constants that are independent of *x*, and which can often be determined from some basic information about the map. Note that, in contrast to the full coding theorem (16), Eq. (17) only provides an upper bound. Nevertheless, a statistical lower bound can be derived [11, 45] showing that most of the probability weight in *P* (*x*) will be close to the bound (17). Interestingly, outputs that are far from the bound can be shown to have inputs (in this case genomes or bonding patterns) that themselves are unusually simple [45].

### B. A Lempel-Ziv based compression approximation for 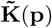

How to best choose a computable complexity measure that approximates the true Kolmogorov complexity is not an easy question to answer, see for example refs. [40, 46–49]) and further discussion in the Supplementary Information of ref. [11]. There is a deep connection between Kolmogorov complexity and compression. For the RNA and GRN, we use a compression method by Lempel and Ziv [50] in which a (binary) string *x* is compressed by looking for patterns (words). The number of words *N_w_*(*x*) forms the basis for this complexity measure that has been a popular choice for approximating Kolmogorov complexity in the literature. In particular, it is thought to work better than many rival methods for shorter strings [51, 52]. In ref. [11] the so-called Lempel-Ziv complexity was defined as follows

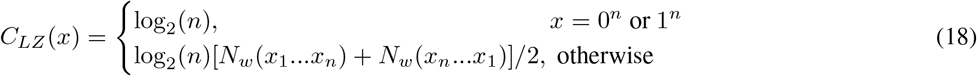

where the simplest strings 0*^n^* and 1*^n^* are separated out because *N_w_*(*x*) which assigns complexity *K* = 1 to the string 0 or 1, but complexity 2 to 0*^n^* or 1*^n^* for *n* ≥ 2, whereas the true Kolmogorov complexity of such a trivial string actually scales as log_2_(*n*), as one only needs to encode *n*. In ref. [11, 24, 45] this complexity measure was empirically shown to work well for a wide diversity of maps, ranging from sets of coupled differential equations, to the RNA SS GP map, to a finite-state transducer, to neural network models for deep learning. This success gives us confidence to use It here for the RNA and the GRN. We note that in the Supplementary Information of ref. [11] we compare a number of other compression based measures for RNA, finding a similar scaling of *P*(*p*) versus 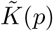 for all of them (note that we use 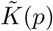 when we are specifically referring to the complexity of some phenotype *p*, whereas 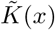 when referring to some general output *x*). Similarly, in ref. [24] we compare a wide range of different complexity measures for a fully connected deep learning model of Boolean functions, finding again that these different measures all correlate with one another, giving us confidence that our results are not that sensitive to the exact method used.

### C. Alternative complexity measures for the polyominoes

**FIG. S13.**
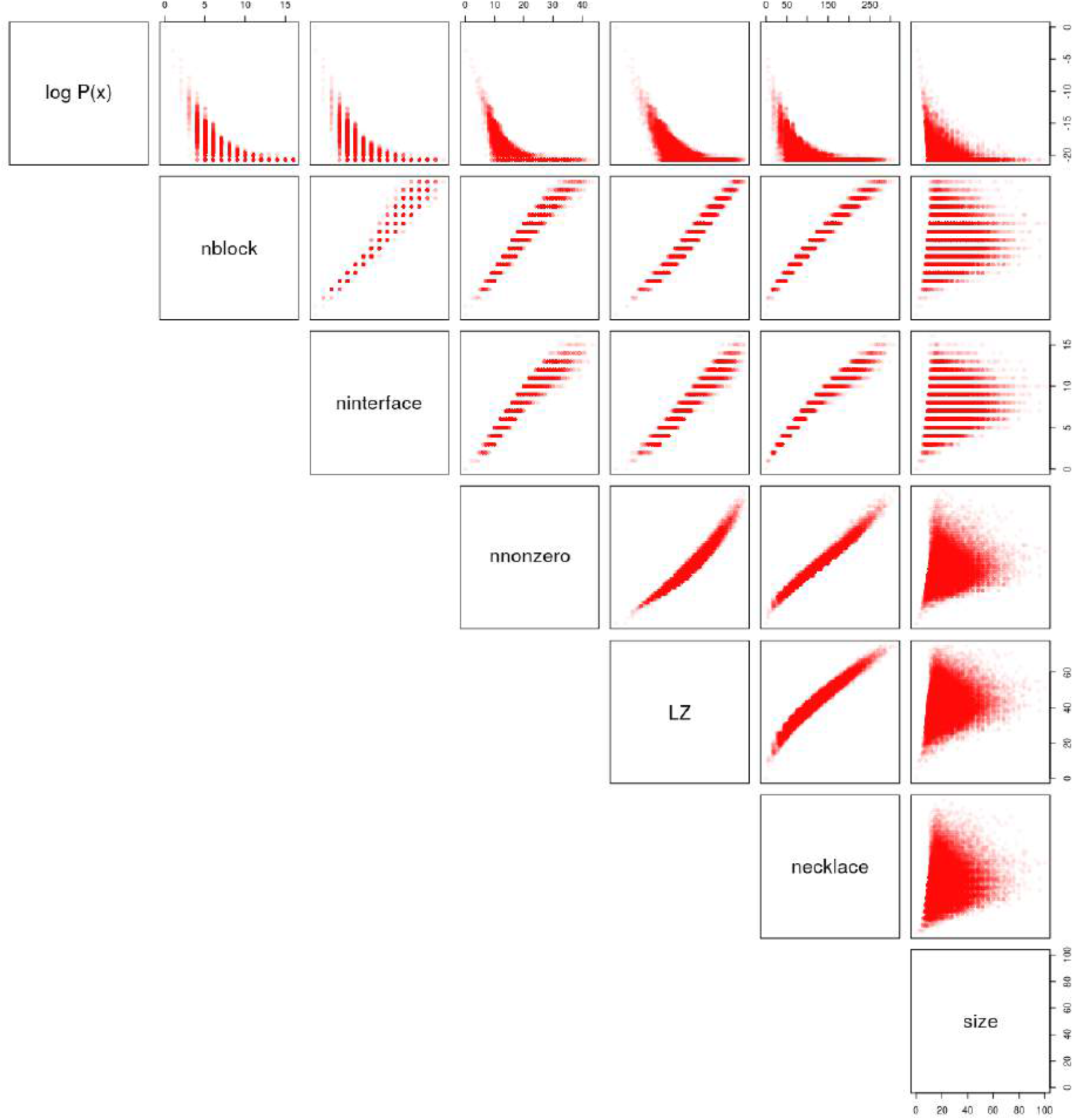
A comparison of different complexity measures for polyominoes for polyomino phenotypes that arise from random sampling of the genotypes in 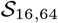 ; The measures are: - nblock = the number *n* of tiles (blocks); - ninterface = the number of interfaces (this measure of complexity 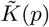 is used in the main text); - nnonzero = the number of non-zero elements of the minimal genome (this directly measures the amount of novel mutations needed); - LZ = our Lempel-Ziv measure from Eq. (18); - necklace = a combinatorial complexity measure from ref. [53] based on necklaces, which can be defined as equivalence classes of strings under rotation.

The complexity measures all generate qualitatively similar *P*(*p*) versus 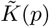 plots, and also correlate relatively well with one another. They all appear to capture some basic properties of the descriptional complexity of the polyominoes. The final row is not a complexity measure, but rather the size of the polyominoes, which does not correlate that well with the complexity measures, as expected, because large structures can be assembled from relatively simple instruction sets. Nevertheless, smaller sizes are more likely to appear than larger sizes, as can be seen in the top right panel. This is because they typically have shorter instruction sets, and so are more likely to appear because fewer evolutionary innovations are needed to make them.

For the polyominoes, we also explored several other complexity measures 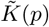 that are all related to the Kolmogorov complexity. These include the number of tiles (blocks), the number of interfaces, the number of non-zero elements of the minimal genome (this directly measures the amount of novel mutations needed), our Lempel-Ziv measure from Eq. (18), and a measure from ref. [53] based on necklaces, which can be defined as equivalence classes of strings under rotation. These measures are all compared in Fig. S13 for the polyomino data from Fig.2 of the main text, that is random sampling of the genotypes. We note that the measures all correlate relatively well with one another supporting our assumption that the exact details of the complexity measure is not critical. They all exhibit the expected exponential scaling between probability and complexity that is predicted by the AIT inspired coding theorem of Eq. (1) in the main text. By contrast, the size of the polyominoes does not correlate well with any of the measures, as expected.

While the more sophisticated Lempel Ziv complexity measure (18) is straightforward to apply to the minimal assembly kit genomes for polyominoes, it is not clear how to use this measure for the protein complexes. In order to compare the polyominoes with the protein complexes, we therefore use the simplified measure that counts the number of interface types. Intuitively, this correlates with the amount of new information that mutations need to supply in order to create a self-assembling polyomino or protein structure, and indeed in Fig. S13 we see that it correlates well with the full Lempel-Ziv complexity measure.

We note that in an important paper [54] Soloveichik and Winfree show that the minimum number of distinct tiles to make a shape scales with the Kolmogorov complexity of that shape. There is a link to our complexity measure because the number of interface types correlates with the number of distinct tile types. Indeed, for the polyominoes, our complexity measure based on the number of tiles correlates closely with the one based on the number of interfaces (see Fig. S13). Further work is needed to flesh out these connections. For the protein complexes, the number of interfaces is a more natural measure than the number of “tiles”.

## S6. SUPPLEMENTARY TEXT FOR BIAS IN OTHER GP MAPS

### 1. Protein tertiary structure

Proteins fold into well-defined three-dimensional structures. Although the folding problem from sequence to structure is highly complex, there are simplified models that can be used to investigate the overall structure of the GP map. A particularly popular model in this regard has been the highly simplified HP lattice model, where folds are represented as self-avoiding walks on a lattice, and the full sequence is reduced to binary alphabet (H stands for hydrophobic and P for polar amino acids) [55]. Despite its heavily coarse-grained nature, this model has produced important biological insights [56]. In particular, a study by Li *et al* [57] showed that for both 2D and 3D models the *NSS*(*p*) distribution was highly biased. For example, in the 3 by 3 by 3 cubic lattice model (with 51704 structures), the number of sequences per structure ranged from ~4000 progressively down to only ~1. Further, it was found that the structures with the largest NSS showed tertiary symmetries and geometrical regularities, similar to the regular and symmetrical forms of natural protein folds, see also [58]. This observation fits within the framework of our observation of the symmetry and simplicity present in large NSS polyominoes, proteins complexes, and RNA structures. From a different perspective, England and Shakhnovich [59] and later Coluzza *et al* [60] have used analytical methods to predict bias in natural protein tertiary structure maps. We note that in this literature, the word “designability” is used to describe how many genotypes (sequences) map to a phenotype (typically a fold).

### 2. Target of rapamycin signalling circuit

Raman and Wagner [61] studied a systems biology model of the target-of-rapamycin (TOR) signalling circuit for budding yeast (*S. cerevisiae*). TOR is a highly conserved protein kinase which controls growth in yeast, fly and mamalian cells. The aim of their study was to see how the topology of the circuit interactions affects the signalling circuit’s behaviour. The circuit’s phenotypes were determined on the basis of the concentration-time trajectories of eight key proteins complexes. Specifically, the model generates a continuous concentration-time profile for each protein, and these were discretised by recording the concentrations at a fixed number of time points. Finally, these trajectories were clustered into similar signalling behaviours using a well known clustering algorithm used for large data sets. The algorithm clustered the (coarse grained) signalling behaviours of key signalling molecules into 286 qualitatively different behaviours, which were then taken as 286 different phenotypes. They then found that this GP map shows very strong bias. For example, the largest *NSS*(*p*) contains 21633 genotypes, or 31% of the total ~70000 genotypes and only 26 phenotypes account for nearly all (82%) of the genotypes. While their method does not obviously allow a complexity measure, it is interesting that it exhibits such strong bias. It would be interesting to attempt to measure the complexity of the phenotypes for this system.

### 3. Boolean threshold models of GRNs

Boolean models of GRNs, where individual genes are simplified to nodes having only an on or off state, and where the temporal dynamics is controlled by a set of Boolean rules for interactions with other nodes (genes), were first introduced by Kauffman [62]. In spite of the great simplification, they have been successfully applied to a wide range of biological phenomena, including segment polarity in *Drosophila* [63], flower development [64], signal transduction in human fibroblasts [65], plant cell signalling [66], and mammalian cortical development [67]. In addition, these models have been used to study more general properties of GRNs, including mutational robustness [68–71] and versions of the concept of evolvability [69–72].

Nevertheless, the state space of Boolean models grows very rapidly with the number of nodes and so it is often hard to achieve good sampling over genotypes. A popular simplification of these models is to use a thresholding rule to determine whether the inputs to a particular node (gene) result in the gene being turned off or on. Such Boolean threshold models (BTNs), which mathematically closely resemble neural networks, have also been successfully used to model GRNs. Examples include GNRs for the mammalian cell cycle [73], the regulation of lymphocyte differentiation [74], and signal transduction in human fibroblasts [65]. Applications of BTNs to the modelling the yeast cell cycle [75–77], have successfully led to predictions of knockout mutant phenotypes [77, 78]. The models have also been used to provide explanations for the designability (redundancy) and robustness of the wild-type phenotype [79–82]. Given their success, it is interesting to think about BTNs as GP maps [9], where the nodes and connections describe the genotypes, and the dynamic behaviour of the model is treated as its phenotype.

In fact, inspired by earlier discoveries of phenotypic bias in HP model lattice proteins [57], Nochomvitz and Li [83] presented evidence for strong bias in BTNs. It has recently been shown that these BTNs also exhibit clear simplicity bias [9], as well as log-linear robustness-frequency behaviour for the mutational robustness, and other properties that are very similar to those observed for other GP maps [13, 15]. Given that we also observed simplicity bias in an ODE system in the main text (where the connectivity is fixed, but the connetions vary in strength), the fact that it is also seen in this quite different and more simplified type of modelling (where the connectivity is varied) suggest that the question of simplicity bias in GRNs may be ripe for further exploration.

### 4. Model of neuron development

Psujek and Beer performed an investigation of phenotypic bias in a computational model of neuronal development [84]. Due to the computational intractability of large neuron networks, the system they developed consists of only three neuron cells, and each pair of neurons *i* and *j* may have a connection arrow (neuronal connection) going from *i* to *j*, from *j* to *i*, or none at all. The phenotype is the connectivity pattern of these neurons, and hence there are 26 = 64 possible phenotypes. The authors observed phenotypic bias on sampling genotypes; interestingly they noted that simpler connection patterns appeared more frequently, which is similar to what we observe in our work with polyominoes, proteins and RNA structures. This behaviour is not surprising in light of more recent papers showing that the perceptron [85] and several architectures for deep neural networks [24] show clear simplicity bias.

